# Connectome analysis reveals brainwide visual processing in *Drosophila*

**DOI:** 10.64898/2026.02.02.700492

**Authors:** Sung Yong Kim, Anmo J Kim

**Affiliations:** Department of Electronic Engineering, Hanyang University, Seoul, South Korea; Department of Biomedical Engineering, Hanyang University, Seoul, South Korea; Department of Artificial Intelligence, Hanyang University, Seoul, South Korea

**Author notes:** Correspondence (A.J.K.).

**Keywords:** *Drosophila* vision, Whole-brain connectome, FAFB-FlyWire, Long-range projection, Brainwide visual processing

## Abstract

Sensory processing relies on an intricate interplay between specialized sensory circuits and the broader brain. Leveraging the whole-brain *Drosophila melanogaster* connectome, we analyzed long-range projections linking the optic lobe (OL)—the principal visual center—reciprocally with the central brain and the contralateral OL to determine how visual features are computed by a brainwide network. By quantifying synaptic polarity and connectivity, we classified the projection neurons into feedforward, feedback, and bilateral groups. Feedforward neurons form an information bottleneck, consolidating diverse visual features from the OL and distributing them through modular pathways for higher-order processing. Feedback neurons densely innervate OL structures, conveying contextual signals in the central brain via precise recurrent loops. Bilateral neurons link local visual circuits across the two OLs, providing a substrate for inter-ocular processing. These findings reveal how early visual pathways interface with brainwide circuits to support diverse brain functions and enable context-dependent flexibility in sensory computation.

**Highlights:**

- Whole-brain connectome analysis identifies three streams of brainwide visual processing
- Feedforward neurons form a bottleneck and distribute features for higher processing
- Feedback neurons modulate early vision via layer-specific recurrent loops
- Bilateral neurons bridge both optic lobes to enable inter-ocular processing

## Introduction

The visual system of *Drosophila melanogaster* is a sophisticated computational engine that calculates a multitude of visual features to facilitate diverse behaviors such as navigation, foraging, courtship, and sleep.^1–8^ The optic lobe (OL) serves as the primary visual center, populated by about 60% of the total neurons in the fly brain and computing various visual features through a hierarchy of parallel processing units.^9,10^ Its role is to analyze a rapidly changing array of light stimuli to extract features relevant to diverse brain functions and distribute them to appropriate neural circuits.

This processing, however, does not occur in isolation; it is intimately coordinated with other brain regions, such as the central brain (CB) and the contralateral OL. The interaction with the CB allows the visual system to dynamically adjust its processing strategies to meet the shifting demands of higher-order functions. For example, the gain of wide-field motion-sensitive cells increases during locomotion, and their activity is suppressed during active flight turns––exemplifying the integration of motor feedback into visual processing.^11–14^ Furthermore, the fly brain performs multi-stage inter-ocular computations that integrate bilateral visual inputs into coherent binocular signals for downstream whole-field visual processing.^15–17^

These brainwide interactions—feature distribution, context-dependent modulation, and inter-ocular comparison—depend on long-range projections that connect the OL with distant regions of the nervous system. Despite the critical importance of these pathways, the signaling architectures of long-range projections in *Drosophila* have not been studied comprehensively.^10,18^

The OL is organized into four major neuropils—the lamina, medulla, lobula, and lobula plate—which maintain a rigorous columnar and retinotopic organization throughout the processing stream. The three proximal neuropils (medulla, lobula and lobula plate) serve as the primary input sources for visual projection neurons (VPNs), which relay visual feature information to the CB. Previous research has characterized the structure and function of a subset of VPNs, alongside visual centrifugal neurons (VCNs)—which convey signals from the CB back to the OL^10,18,19^—and a few types of bilateral neurons that link the two OLs.^20–22^

The recent completion of the synapse-level wiring map of the entire *Drosophila* brain (FAFB-FlyWire) presents a transformative opportunity to resolve these circuits.^10,19,23–25^ While "projectomes" in larger mammals, such as primates and mice, have mapped long-range connectivity at the mesoscale, they lack the resolution to identify specific synaptic partners across the entire population.^26,27^ The *Drosophila* connectome now allows us to trace the precise "input-output" logic of long-range projections, revealing how information flows to and from visual pathways with synaptic specificity.^19,26^

In this study, we performed comprehensive analyses of projection neurons that connect the OL with other brain regions. Specifically, based on their connectivity profiles, we classified these neurons into three functional categories: feedforward (FF), feedback (FB), and bilateral (BL) (Figure 1A). To elucidate their respective roles in visual processing, we examined the projection profiles of FF neurons within the CB and identified functional modules defined by shared downstream targets. We next characterized the innervation patterns of FB neurons in the OL, uncovering a layer-specific recurrent loop architecture that likely modulates early visual processing according to various higher-order signals in the CB. Finally, we mapped the receptive and projective fields of BL neurons and analyzed their connectivity to predict their functional roles in combining visual information across the two OLs.

**Figure 1.**
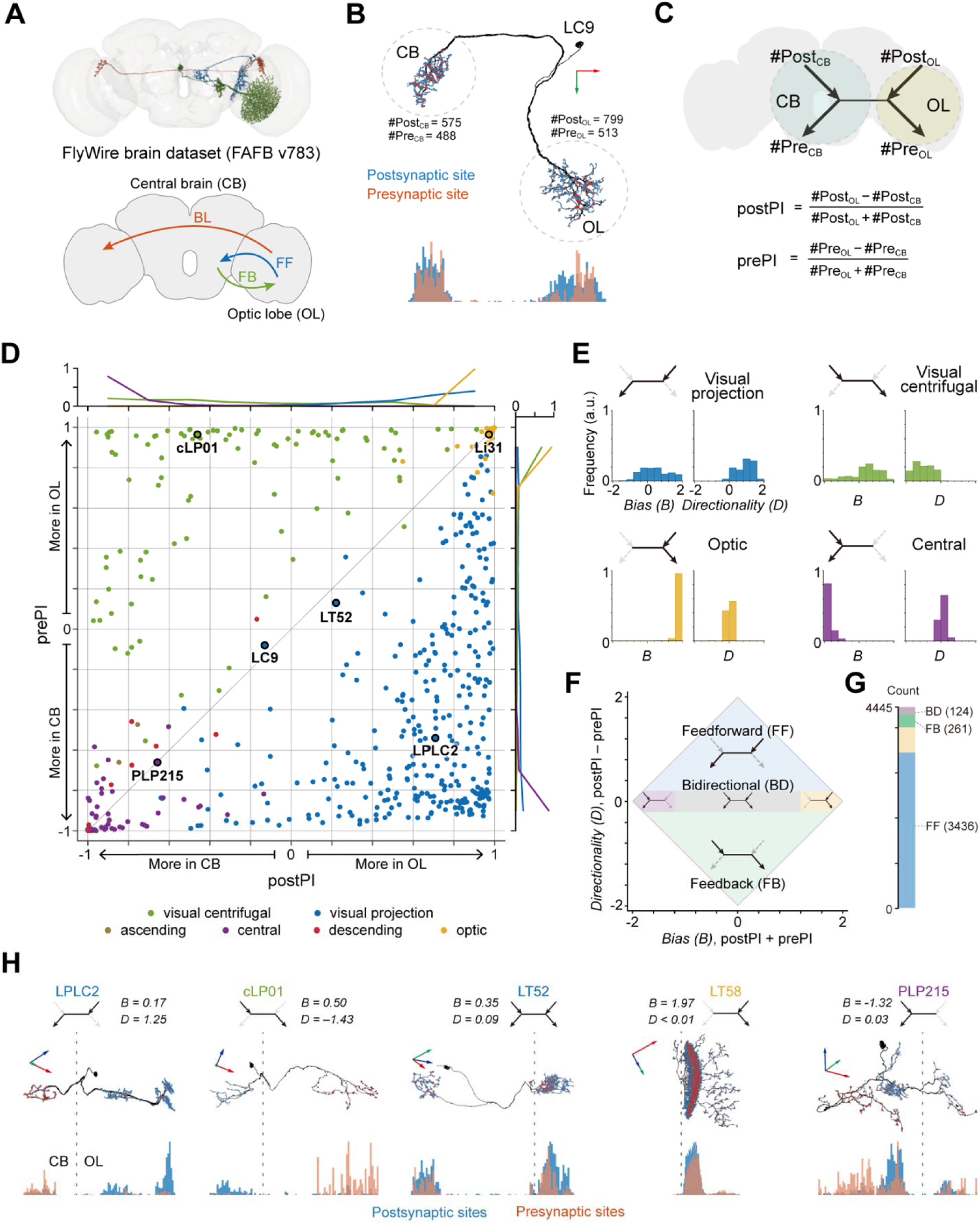
Classification of projection neurons by the distribution of synapses. **(A)** Representative morphologies of one feedforward (FF), one feedback (FB), and one bilateral (BL) neuron in the FAFB-FlyWire connectome (top), and a schematic illustrating the corresponding directions of information flow (bottom). **(B)** Distribution of pre- and post-synapses within a sample LC9 neuron. The histograms at the bottom depict the density of pre- and post-synaptic sites in the optic lobe (OL) and the central brain (CB), with the x-axis representing physical distance in the brain. **(C)** Schematic defining polarity indices for presynaptic (prePI) and postsynaptic (postPI) sites between the OL and CB. **(D)** Two-dimensional scatter plot of postPI and prePI values for all cell types linking the right OL and CB. Each dot represents a cell type and is color-coded by superclass. Histograms of PI values for each cell type are shown along the top and right margins. **(E)** Histograms of spatial bias (*B =* postPI + prePI) and directionality (*D =* postPI - prePI) across four superclasses: visual projection, visual centrifugal, optic, and central neurons. **(F)** Two-dimensional depiction of *B* (spatial bias) and *D* (directionality), along with cell type classification criteria for FF, BD, and FB neurons. **(G)** Stacked bar plot showing the number of neurons in each region defined in (F). Colors correspond to those in (F). Labels indicate FF, BD, and FB neuron counts. **(H)** Example neurons from each class: FF, FB, BD, optic, and central (left to right). Synapse distribution histograms are shown below as in (B). The dotted line indicates the boundary between the OL and the CB. Red, green, and blue arrows denote the x-, y-, and z-axes in the connectome, respectively, which are approximately aligned with the brain’s lateral, dorsoventral, and anteroposterior axes.

## Results

### Classification of projection neurons based on synaptic distribution

The long-range projections linking the OL and the CB are primary conduits for visual information. However, their signaling direction—FF or FB—often cannot be determined from morphology alone because *Drosophila* neurons often feature mixed pre- and post-synaptic sites within the same compartment, such as LC9 neurons (Figure 1B).^28^

To determine the signaling direction, we quantified the distribution of synapses to define polarity indices (PIs) for postsynaptic (postPI) and presynaptic (prePI) sites for all the neurons that link the right OL and the CB (Methods; Figure 1C). postPI measures the distribution of postsynaptic sites between the OL and the CB; the value of 1 indicates that all postsynaptic sites are located in the OL, while -1 indicates all sites are within the CB. A value of 0 denotes an even distribution across both regions. prePI is defined in the same way, but for presynaptic sites. Analysis of (postPI, prePI) values for individual cell types revealed a strong bimodal distribution (Figure 1D).

Neurons previously annotated as VPNs clustered in the lower-right area below the diagonal (postPI > prePI), indicating input dominance in the OL and output dominance in the CB. Conversely, VCNs occupied the upper-left area (prePI > postPI). We noted that some neurons previously annotated as intrinsic to the OL (termed ‘optic’ neurons in FlyWire) were included in our analysis; these clustered in the upper-right corner (both indices ≈ 1) due to a small fraction of their synapses being located in the CB. Similarly, neurons previously labeled as intrinsic to the CB (’central’ neurons in FlyWire) clustered in the lower-left corner (both indices ≈ -1).

We further refined this analysis by defining two composite metrics from the PIs to capture higher-order properties of synaptic distribution. The spatial bias (*B*) was defined as the sum of the two indices (postPI + prePI); values near 2 or -2 indicate neurons restricted to the OL or CB, respectively, while values near 0 indicate a balanced spatial distribution. The directionality index (*D*) was defined as the difference between the two PIs (e.g., postPI - prePI). Values near 2 indicate FF signaling (i.e., OL-to-CB), values near -2 indicate FB signaling (i.e., CB-to-OL), and values near 0 suggest neutral directionality. When we analyzed neurons grouped by major "superclasses" (per FlyWire annotations^10^) using these metrics, we found that different superclasses exhibited distinct combinations of directionality and bias indices (Figure 1E). For example, VPNs exhibited a broad range of bias leaning slightly positive, paired with strictly positive directionality.

Using these metrics, we re-classified the projection neurons into distinct functional groups (Figures 1F and 1G). We identified FF neurons as those with *D* ≥ 0.2 (3,436 cells; e.g., LPLC2 in Figure 1H), and FB neurons as those with *D* ≤ -0.2 (261 cells; e.g., cLP01 in Figure 1H). Over 99% of FF neurons were previously annotated as VPNs, and 95% of FB neurons were previously annotated as VCNs. Finally, a small subset of neurons exhibiting both a balanced spatial distribution and near-zero directionality were classified as bidirectional (BD) neurons (124 cells; |*D|* < 0.2 and |*B|* < 1.2; e.g., LT52 in Figure 1H). Among neurons with weak directionality (|*D| < 0.2*), we excluded neurons with |*B|* ≥ 1.2 as their signaling is mostly limited in either the OL or CB. Additional analysis for the neurons connecting the left OL with the CB exhibited similar (postPI, prePI) values for the same type of cells (Figure S1), confirming that our method reliably estimates signaling direction regardless of brain laterality. In the following, we focused our analyses on the connectivity of these FF, FB, and BD neurons.

### Visual features converge in FF neurons and are reintegrated in the CB

FF neurons represent the primary pathway for relaying visual information from the OL to the CB (Figure 2A, top). The major FF cell types include lobula columnar (LC), medulla tangential (MTe), and lobula plate-lobula columnar (LLPC) neurons (Figure 2A, bottom). A small subset of FF neurons (n = 40) extends neurites to the contralateral OL, but we included them as FF cells because they exhibited significant connectivity from the right OL to the CB (Methods).

**Figure 2.**
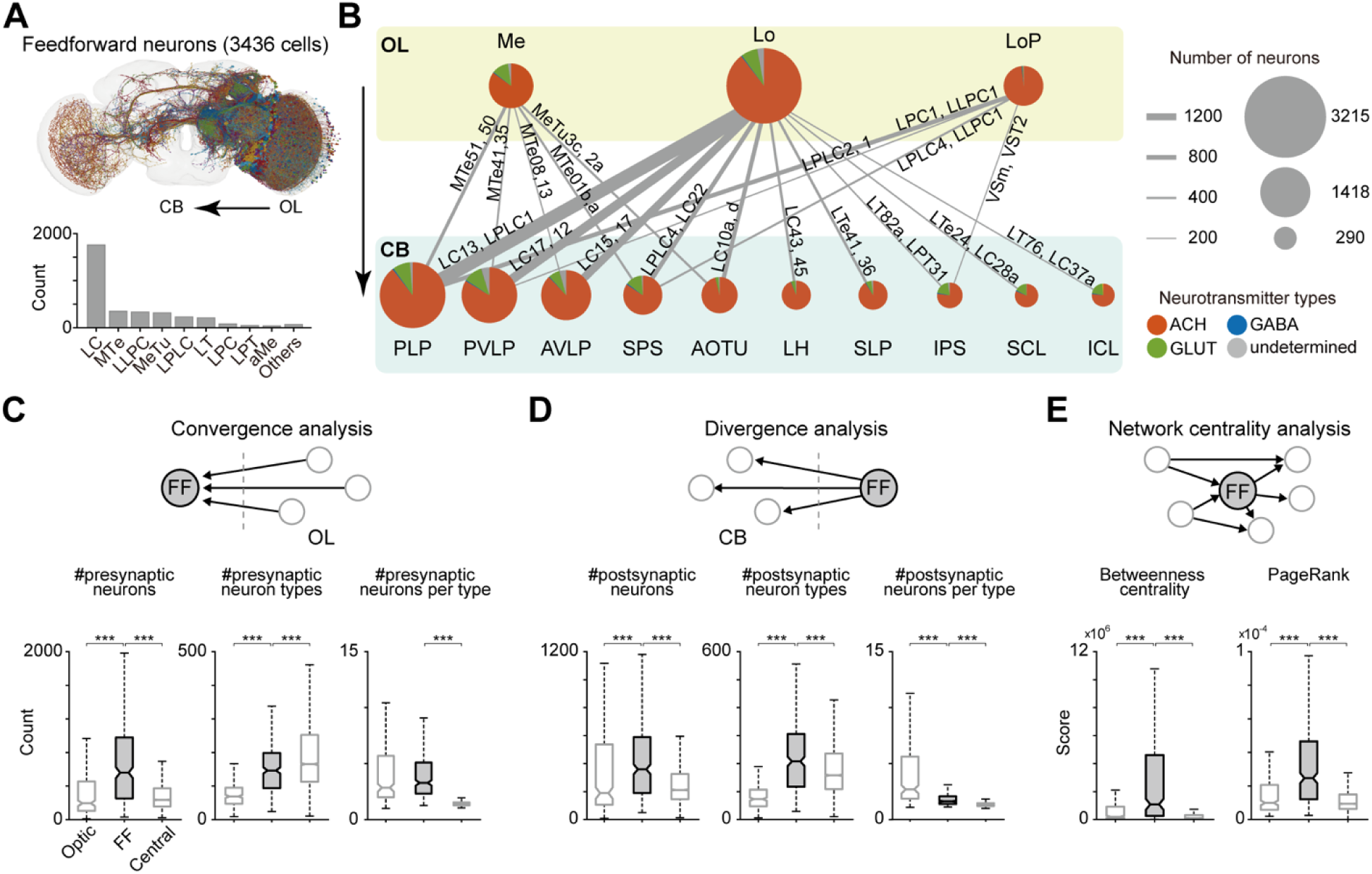
Characterization of convergence and divergence properties of FF neurons. **(A)** Morphologies of all FF neurons projecting from the right OL to the CB (top), and cell counts for each FF neuron type (bottom). **(B)** Schematic of neuropil-level connectivity for FF neurons. The diameter of each circle and the width of each link are proportional to the number of cells in each group. The pie chart shows neurotransmitter identity. Cell-type names on links indicate the two neuron types with the highest synapse count for each connection. **(C)** Input convergence analysis. Boxplots show the number of presynaptic neurons across optic, FF, and central classes (left), the number of presynaptic cell types (middle), and the number of presynaptic neurons per presynaptic cell type (right). **(D)** Output divergence analysis. Boxplots show the number of postsynaptic neurons across optic, FF, and central classes (left), the number of postsynaptic cell types (middle), and the number of postsynaptic neurons per postsynaptic cell type (right). **(E)** Network centrality analysis. Boxplots show betweenness centrality (left) and PageRank (right) across optic, FF, and central classes. Statistics (C–E): Mann–Whitney U test with Bonferroni-corrected α = 0.0167; asterisks denote significance (*p < 0.0167; ***p < 0.0001).

To characterize how these neurons transmit visual features, we analyzed the neuropils housing their input and output synapses (Figure 2B). The lobula provided the largest source of input to FF neurons, followed by the medulla and the lobula plate. Conversely, FF outputs primarily targeted CB neuropils associated with higher-order visual processing, including the anterior optic tubercle (AOTU), anterior ventrolateral protocerebrum (AVLP), posterior lateral protocerebrum (PLP), and posterior ventrolateral protocerebrum (PVLP). Specifically, medulla-originating FF neurons most frequently projected to the AOTU,^29^ whereas those originating in the lobula and lobula plate most often targeted the PLP.

We also found that FF neurons route substantial input to CB regions known primarily for non-visual functions. A subset of the lobula FF neurons projects to the superior posterior slope (SPS) and inferior posterior slope (IPS), which serve as major input sources for descending neurons involved in motor control.^30,31^ Furthermore, FF neurons from the lobula innervate regions implicated in circadian rhythms—such as the superior lateral protocerebrum (SLP), superior clamp (SCL), and inferior clamp (ICL)^32^—as well as the lateral horn (LH), a center traditionally associated with innate olfactory behaviors.^33^ These wiring patterns demonstrate that visual features extracted in the OL are broadly distributed to central circuits coordinating diverse brain functions.

FF neurons gain their visual selectivity by integrating convergent inputs from the presynaptic optic neurons. We found that FF neurons receive input from a significantly larger number of presynaptic partners compared to optic and central neurons (Figure 2C, left). This high degree of convergence could arise from two distinct wiring strategies: the integration of diverse visual features (pooling different cell types) or the integration of the same features arising in different visual columns (pooling many neurons of the same type). Our analysis supports both mechanisms. Namely, FF neurons sample from a greater variety of presynaptic cell types than optic neurons, likely contributing to the emergence of novel feature selectivity (Figure 2C, middle). Furthermore, the number of presynaptic neurons per type is significantly higher than central neurons, supporting a high degree of spatial integration that may help expand receptive fields and enhance signal robustness (Figure 2C, right).^34–36^

We next examined how these integrated visual features are disseminated to the CB. FF neurons synapse onto a significantly larger number of postsynaptic cells and cell types than both optic and central neurons (Figure 2D, left and middle). This high degree of output divergence supports the parallel recruitment of distinct functional pathways, allowing a single visual feature to simultaneously update systems for different brain functions.^37–40^ However, the average number of postsynaptic cells per type is lower for FF neurons than for optic neurons (Figure 2D, right). This likely reflects a fundamental difference in circuit architecture: whereas optic neurons broadcast redundantly to many neurons of the same type, FF neurons prioritize distributing information across a diverse array of functionally distinct targets.^6,38,39,41–43^

Collectively, these findings identify the FF population as a structural bottleneck in the visual processing hierarchy. As manifested by high input convergence and high output divergence, these neurons consolidate spatially and functionally diverse signals from the OL before distributing them widely across the CB. To test this “bottleneck” role, we computed centrality measures: betweenness captures shortest-path bridging,^44^ whereas PageRank captures neighbor-weighted global influence.^45^ FF neurons exhibited significantly higher betweenness centrality and PageRank scores than either optic or central neurons (Figure 2E). These metrics confirm that FF neurons act as critical, influential nodes that facilitate the efficient compression and distribution of information across the brainwide visual network.

### Clustering analysis reveals the functional modules of FF neurons

FF neurons distribute visual features to diverse neural circuits in the CB. A CB neuron downstream of the FF population may either directly utilize signals from a FF cell type or integrate inputs from multiple types to generate more complex representations. To distinguish between these two possibilities, we quantified the number of distinct FF cell types providing direct synaptic input to individual CB neurons. Of the 7,552 CB neurons postsynaptic to the FF population, 58% received inputs from more than one FF cell type (Figure 3A). This widespread convergence suggests that a majority of CB neurons serve as sites for feature integration. However, it remains unresolved whether such convergence primarily reflects the pooling of similarly tuned FF neurons to enhance signaling robustness, or the synthesis of novel, higher-order features through the integration of distinct visual channels.

**Figure 3.**
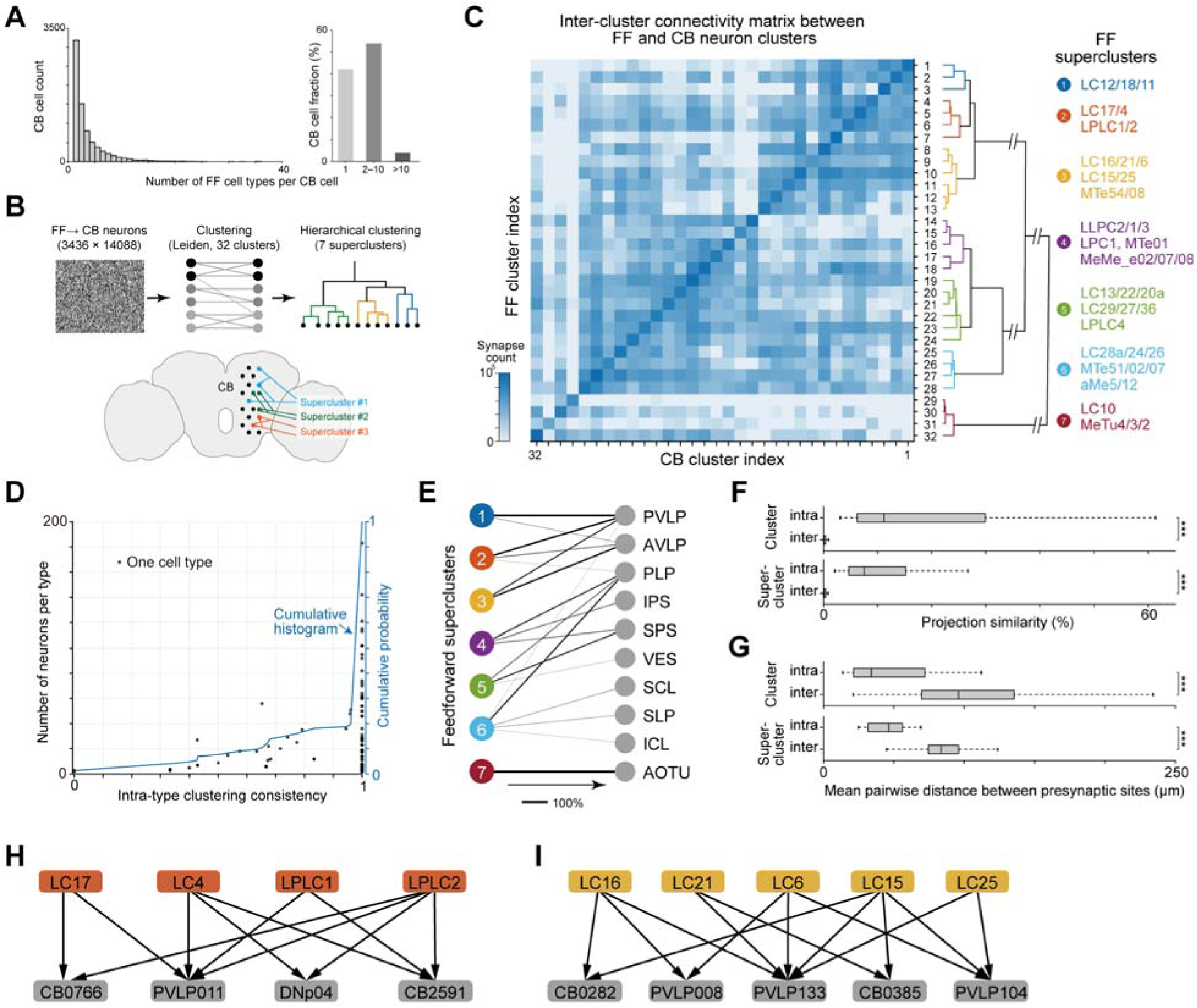
Modular projection patterns of FF neurons onto CB circuits. **(A)** Histogram showing the distribution of the number of presynaptic FF input types per postsynaptic neuron in the CB (left). Summary bar plots show the fraction of postsynaptic CB neurons in three bins of presynaptic FF input-type counts (1, 2–10, and >10) (right). Only connections with synaptic weight ≥ 5 were included in the analysis. **(B)** Schematics of the analysis workflow (top) and the FF clusters (bottom). **(C)** Connectivity matrix from FF clusters (rows) to postsynaptic target clusters (columns) (left). Dendrogram showing distances between FF clusters based on their projection patterns (right). Colors in the dendrogram indicate superclusters—groups of clusters whose dendrogram distance falls below a threshold—and representative neuron types within each supercluster are labeled on the right. **(D)** Clustering consistency across neuron types with at least two neurons per type. Each dot represents a neuron type; the x-axis shows the fraction of within-type neuron pairs assigned to the same cluster, and the left y-axis shows the number of neurons in that type. The cumulative distribution of clustering consistency is plotted as a blue line (right y-axis). **(E)** Neuropil-level projection profiles of superclusters. Edge thickness denotes the fraction of output synapses from each supercluster to each neuropil. Supercluster colors correspond to those shown in (B). **(F)** Box plots of FF projection similarity, comparing intra-versus inter-group pairs for clusters (top) and superclusters (bottom), quantified by the weighted Jaccard index. **(G)** Box plots of mean pairwise distance between presynaptic sites, comparing intra-versus inter-group pairs for clusters (top) and superclusters (bottom). **(H)** Convergence circuits in the CB for FF neurons in supercluster #2. **(I)** Convergence circuits in the CB for FF neurons in supercluster #3. In (H) and (I), FF neuron colors match those in (C). Statistics (F–G): Mann–Whitney U test; asterisks denote significance (***p < 0.0001).

To investigate the logic of this integration, we sought to identify functional modules of FF neurons based on their downstream connectivity profiles. To identify FF modules, we applied a modularity-based community detection method to a bipartite graph between FF neurons and their CB postsynaptic partners (Methods; Figure 3B).^46,47^ This process assigned each FF neuron to a specific module whose members shared significantly more postsynaptic partners with each other than with neurons in other communities. Through this analysis, we identified 32 distinct clusters of FF neurons (Figure 3C; Table S1). Notably, this method reliably grouped neurons of the same annotated type into the same clusters, despite significant variability in connectivity profiles across individual neurons of the same type (Figure 3D; Table S1).^48,49^ To further characterize the relationship between these clusters, we quantified the similarity between individual clusters via hierarchical clustering (Methods; Figure 3C).^47,50^ By applying a threshold to the resulting dendrogram, we eventually identified seven "superclusters" of FF neurons (Figures 3C, 3E, and S2), each representing neurons sharing broad targeting motifs (Figure 3E). For example, superclusters #1, #2, and #3 primarily innervated the PVLP and AVLP, whereas superclusters #4 and #5 targeted the PLP and SPS. Superclusters #6 targeted the SCL, SLP, and ICL, while #7 projected exclusively to the AOTU.

Our analysis confirmed that neurons within the same cluster or supercluster share a significantly larger fraction of postsynaptic partners (Figure 3F). For individual clusters, the average projection similarity was 19.8 %, substantially higher than the 0.4 % observed between neurons from different clusters. At the supercluster level, intra-group similarity averaged 10.5 %.

Furthermore, because neurons with high projection similarity tend to target the same CB regions, the physical distance between their output synapses was also significantly shorter for neurons within the same cluster or supercluster (Figure 3G).

At the supercluster level, we observed that neurons with similar feature selectivity converge onto shared downstream targets. This is exemplified by supercluster #2 (Figure 3H; Table S1), which groups LC17, LC14, LPLC1, and LPLC2, all of which were previously shown to respond strongly to looming objects.^51,52^ While the lack of visual tuning data for the majority of FF neurons precludes a comprehensive quantitative analysis, we found consistent convergence of similarly tuned FF neurons onto common downstream targets across multiple superclusters (e.g., supercluster #1 for translating spots and supercluster #5 for looming discs). Such convergence may enhance the robustness of the visual feature representations used by CB neurons.^51,53^ Conversely, we also identified instances where FF neurons with distinct visual tuning converge. For instance, supercluster #3 comprises a diverse array of types, including LC6 and LC16 (looming-sensitive), LC21 (spot-sensitive), LC15 (bar-sensitive), and LC25 (grating-sensitive) (Figure 3I; Table S1).^51,54^ Such heterogeneous convergence may facilitate the synthesis of novel, higher-order visual features within downstream circuits.^55–57^

Together, our clustering analysis revealed organized functional groups of FF neurons defined by shared postsynaptic targets. While the full functional implications of these clusters remain to be determined, our results suggest that these modules represent specific visual channels that exert coordinated influence over downstream central and motor circuits.

### FB neurons convey diverse CB signals to the OL, via a precise recurrent loop

FB signaling enables active sensing by modulating sensory processing according to an organism’s internal states or sensory variables in the CB. In *Drosophila* vision, previous work has implicated FB neurons at multiple levels of the visual pathway in functions such as detecting directional motion,^58^ filtering low-frequency signals,^59^ and suppressing self-generated visual inputs during flight turns and grooming.^11,13^

Based on the postPI-prePI analysis (Figure 1), we identified 261 FB neurons that route information from the CB to the right OL (Methods; Figure 4A, top). Specifically, the FB cell group comprised 247 VCNs alongside a small population of non-VCNs (two ascending, three descending, three central, and six optic neurons). The most prevalent types of FB neurons were cLP (n = 96), cL (n = 46), and cM (n = 42) (Figure 4A, bottom).

**Figure 4.**
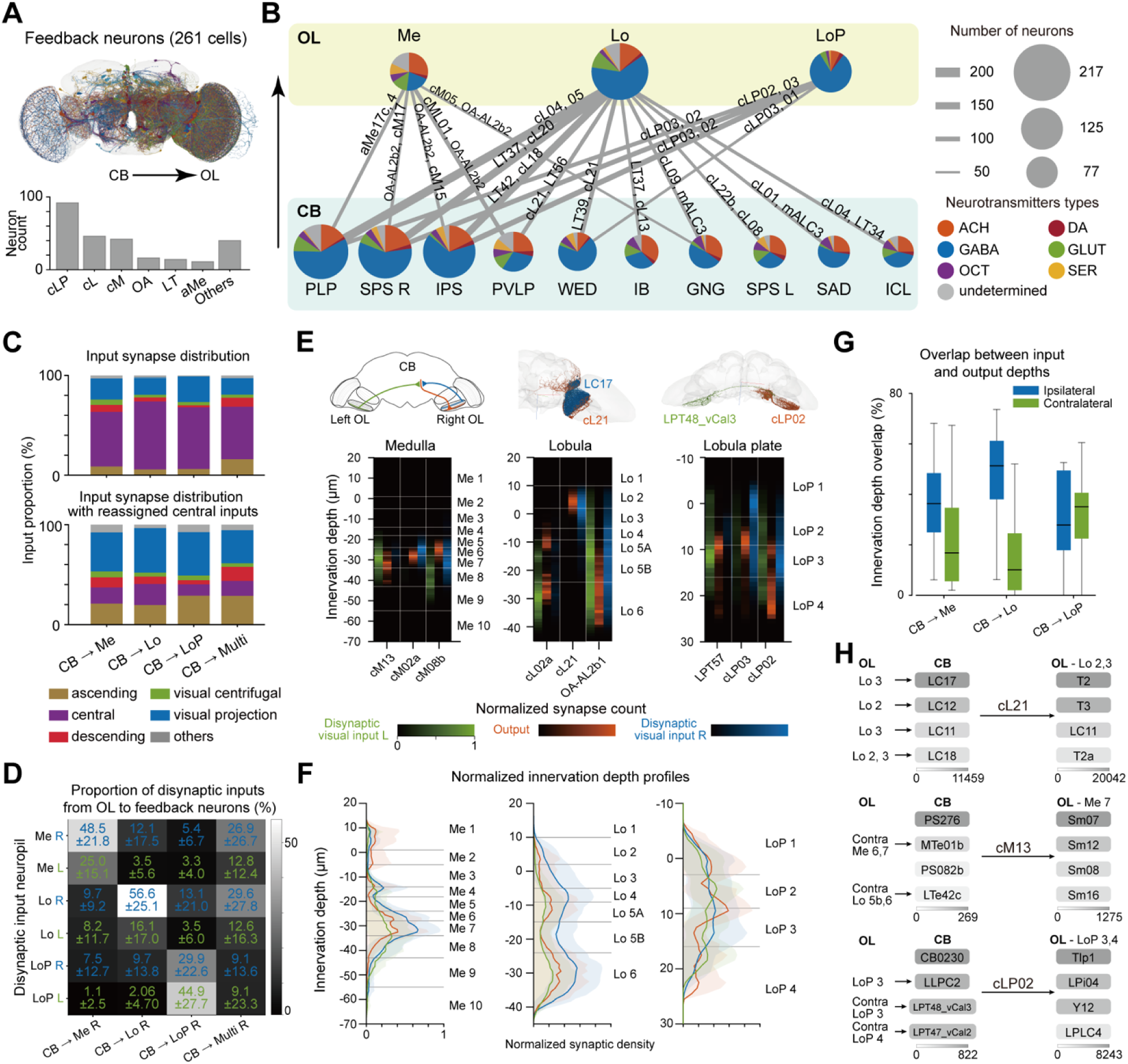
FB neurons integrate diverse CB signals via precise reciprocal loops. **(A)** Morphologies of all FB neurons projecting from the CB to the right OL (top), and cell counts for each FB neuron type (bottom). **(B)** Schematic of neuropil-level connectivity for all FB neurons. The diameter of each circle and the width of each link are proportional to the cell count in each group. Pie charts depict neurotransmitter distributions. Cell-type names on links indicate the two neuron types with the highest synapse counts. **(C)** Superclass distribution of neurons presynaptic to four groups of FB neurons (top). The superclass distribution is replotted after reassigning central neurons according to their input profiles (bottom, Methods). **(D)** Proportion of disynaptic inputs from different OL neuropils. Values are mean ± s.d. **(E)** Innervation-depth profiles of disynaptic visual inputs and outputs for sample FB neurons (bottom), corresponding diagrams (top left), and sample neuron images (top middle and right). Depth profiles are color-coded green for contralateral input, blue for ipsilateral input, and red for output sites. **(F)** Innervation-depth distributions of disynaptic visual inputs and outputs for the three neuropils. Solid lines indicate the mean; shaded regions indicate ± s.d. **(G)** Sum of the minimum overlaps between depth-binned density profiles of output sites and disynaptic inputs. Green indicates overlap with contralateral input; blue indicates ipsilateral. **(H)** Connectivity diagrams for three representative FB neurons. For each side of the neuron, the top four connected partners are shown. Box color intensity corresponds to connection strength.

We found that the lobula received the largest share of FB outputs, followed by the lobula plate and the medulla (Figure 4B). Notably, the neuropils identified as the principal targets of FF neurons—the PLP, PVLP, IPS, SPS, and ICL—also serve as the primary sources for FB neurons returning to the OL, suggesting the existence of recurrent circuit motifs involved in the visual processing. Additionally, the wedge (WED), the gnathal ganglion (GNG), the saddle (SAD), and the inferior bridge (IB) contributed significant inputs to some FB neurons. These pathways likely provide diverse internal state and contextual signals via FB neurons to dynamically modulate visual processing within the OL.^31,60–62^

To understand the nature of signals conveyed by FB neurons, we analyzed the superclass of presynaptic partners within the CB (Figure 4C, top). Among these superclasses, central neurons contributed the largest fraction of inputs, followed by VPNs. However, because the central neuron superclass encompasses diverse functional populations, we further resolved their functional identity by reassigning them based on the superclasses of their own presynaptic partners (Methods; Figure 4C, bottom). This second-order analysis revealed that the majority of inputs to FB neurons are ultimately attributable to VPNs (∼41%), followed by ascending neurons (Methods; Figure 4C, bottom and Figure S3A ).

FB neurons receiving inputs from VPNs form a recurrent loop by pooling visual signals disynaptically from the OL and subsequently returning them to it. To determine the spatial relationship between the origin and target regions for different FB populations, we compared the distribution of their disynaptic visual inputs against their output sites within the OL (Methods; Figures 4D and S3B). We observed a significant overlap between the origin and target neuropils. Specifically, FB neurons projecting to the right medulla received 48.5 ± 21.8% of their disynaptic visual input from the ipsilateral medulla. Similarly, neurons targeting the right lobula received their largest share of input from the ipsilateral lobula (56.6 ± 25.1%). These fractions were significantly larger than those from the contralateral OL neuropils. In contrast, for neurons targeting the right lobula plate, contralateral inputs exceeded ipsilateral ones (contra 44.9 ± 27.7% vs. ipsi 29.9 ± 22.6%), suggesting that the FB pathways to the lobula plate may contribute to bilateral visual integration.

To understand the nature of this reciprocity in further detail, we quantified how disynaptic input and monosynaptic outputs for individual FB neurons are organized in OL layers, which are known to be associated with distinct visual computations (Methods; Figures 4E, S3C, and S3D).^52,54,63–68^ Sample FB neurons with a high degree of input/output alignment include neurons receiving contralateral inputs (e.g., cM13 in the medulla; cL02a in the lobula; LPT57 in the lobula plate), those receiving ipsilateral inputs (e.g., cM02a in the medulla; cL21 in the lobula; cLP03 in the lobula plate), and those integrating inputs from both sides (e.g., cM08b in the medulla; OA-AL2b1 in the lobula; cLP02 in the lobula plate) (Figure 4E).

We also analyzed the density of input and output synapses across layers. We found that FB neurons exhibited stereotyped innervation patterns for both disynaptic input and output synapses (Figure 4F). For example, medulla-projecting FB neurons exhibited the highest density in medulla layer 7 for both disynaptic inputs and presynaptic sites (Figure 4F, left).^63,69^ Lobula-projecting neurons showed the highest density of both sites in lobula layers 4 and 6 (Figure 4F, middle). In contrast, the disynaptic input and presynaptic sites of FB neurons projecting to the lobula plate were broadly distributed across layers (Figure 4F, right). When we measured the strength of reciprocity by summing the minimum overlap between input and output density profiles across depth bins, we found a high degree of overlap in all ipsilateral neuropils (Figure 4G and S3E). Conversely, we found that contralateral overlap is comparable to that of ipsilateral loops only in the lobula plate.

For some FB neurons, the strong reciprocity was evident even when analyzing individual, directly connected cell types (Figure 4H, top). For example, cL21 neurons pool inputs disynaptically from several LC neurons (LC17, LC12, LC11, and LC18) and return to them either directly (LC11) or disynaptically via optic neurons (e.g., LC11, LC17, and LC18 are postsynaptic to T2 or T3). For other FB neurons, however, cell-type-level reciprocity was not apparent. For instance, while cM13 and cLP02 cells formed precise recurrent loops in their innervation layers, their specific upstream and downstream cell types appeared disjoint (Figure 4H, middle and bottom). Together, these findings suggest that while some FB neurons act directly upon specific feature-encoding VPNs, the majority of FB circuits interact indirectly within a specific layer. Such recurrent architecture in the early visual system may facilitate the dynamic modulation of feature sensitivity within or across visual processing pathways.^59,70–72^

### A small subset of projection neurons exhibit directionally neutral synaptic polarity

Based on the spatial distribution of synapses, we defined BD neurons as those that exhibited highly balanced expression of synapses in the right OL and the CB, with a low directionality (Figures 1F, 5A, and 5B). For a representative set of BD neurons (LC9, LT43, and LT52), their mean polarity indices (postPI, prePI) were both near zero, positioning them midway between FF and FB neurons (Figures 5A-D). The postPI for the entire BD neuron population (11 cell types, 124 cells) differed significantly from that of FF neurons, whereas their prePI differed significantly from either prePI or postPI (Figures 5B-D).

**Figure 5.**
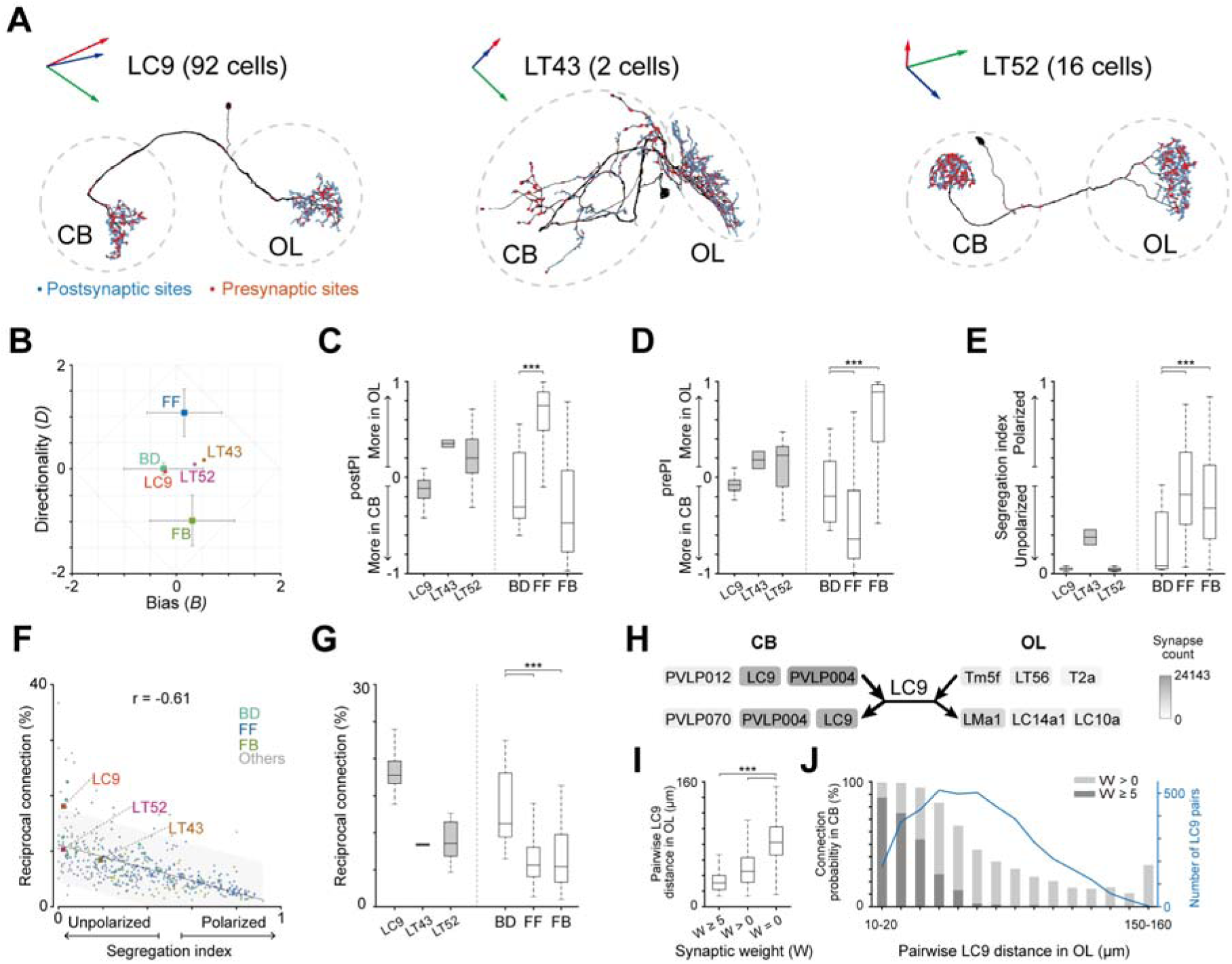
A small subset of projection neurons exhibit bidirectional connectivity. **(A)** Morphologies of representative bidirectional (BD) neurons and the locations of their pre- and postsynaptic sites. Red, green, and blue arrows represent the x-, y-, and z-axes in the connectome, respectively, which are approximately aligned with the brain’s lateral, dorsoventral, and anteroposterior axes. **(B)** Directionality (*D*) and spatial bias (*B*) for BD, FF, and FB neurons. Each point denotes the mean, and error bars indicate ±1 s.d. The BD neurons shown in (A) are highlighted. **(C)** Box plots of postPI at the cell level for representative BD neurons (left of the dotted line) and at the type level for BD, FF, and FB (right of the dotted line). **(D)** Same as (C), but for prePI. **(E)** Box plots of segregation index at the cell level for representative BD neurons (left) and at the type level for BD, FF, and FB (right). **(F)** Scatter plot relating the segregation index to reciprocal connectivity, showing a negative correlation (Pearson’s r = −0.61). Each dot represents one neuron type. The gray shaded area indicates the 95% prediction interval of the linear fit. The BD neurons shown in (A) are highlighted. **(G)** Box plots of reciprocal connectivity fraction, for sample cells (left) and for cell types (right). **(H)** Connectivity diagrams showing, for LC9, the three top presynaptic and postsynaptic partners in the OL and CB. Box color indicates the synapse count between the partner and the LC9 neuron (see colorbar). **(I)** Box plots of pairwise distances between LC9 neurons in the OL as a function of synaptic weight (*W*). *W* = 0 indicates no detected connection, whereas *W* > 0 indicates the presence of at least one synaptic connection. *W* ≥ 5 denotes LC9 pairs whose total reciprocal synapse count (LC9→LC9 + LC9←LC9) is ≥ 5. **(J)** Graph showing the relationship between pairwise LC9 distance in the OL and (i) the number of LC9 pairs (right y-axis) and (ii) the probability of synaptic connectivity (left y-axis), quantified separately for connections with *W* > 0 and for stronger connections with *W* ≥ 5. Statistics (C–E, G): Mann–Whitney U test with Bonferroni-corrected α = 0.0167; asterisks denote significance (*p < 0.0167; ***p < 0.0001).

We also measured the segregation index of these neurons to quantify the degree of spatial intermixing of synapses within each neuron (Figure 5E).^73^ BD neurons consistently exhibited near-zero segregation indices. Higher segregation indices reflect greater spatial separation between pre- and postsynaptic sites within a neuron, reducing the likelihood that partner neurons form reciprocal connections. We tested this by quantifying reciprocity for BD neuron partners (Methods). Analysis revealed a significant negative correlation between reciprocity and the segregation index across all projection neurons (Figure 5F). Notably, even for BD neurons—which exhibited the highest reciprocity—the median reciprocal-connection ratio remained below 20% (Figure 5G). This low degree of cell-level reciprocity underscores the precise and predominantly unidirectional nature of synaptic targeting, even within BD neurons.

Analysis of the connectivity between representative BD cell types and their primary partners revealed that reciprocal connections are localized predominantly within the CB. For example, LC9 cells establish robust reciprocal motifs with PVLP004 neurons in the CB (Figure 5H). Furthermore, we observed prominent direct lateral connections among LC9 neurons (Figure 5H). Notably, when two LC9 neurons were connected in the CB, their pairwise distance within the OL tended to be short (Figure 5I). Conversely, LC9 pairs that were closer together in the OL were more likely to be synaptically connected (Figure 5J). This may suggest that LC9 neurons may expand their receptive field by connecting with other LC9 neurons that have a neighboring receptive field.

### Diverse BL neurons bridge the visual circuits of both OLs

Many projection neurons form direct connections between the right and left OLs, providing a key neural substrate for bilateral visual integration at the early sensory pathway. We first calculated the polarity indices (postPI, prePI) for neurons annotated as right-sided by FlyWire, based on the number of synapses in the right versus left OL (Figure 6A). In the (postPI, prePI) space, most neurons were positioned in the lower-right area below the diagonal (postPI > prePI), suggesting that input synapses are predominantly located in the right OL and output synapses in the left OL (Figure 6B).

**Figure 6.**
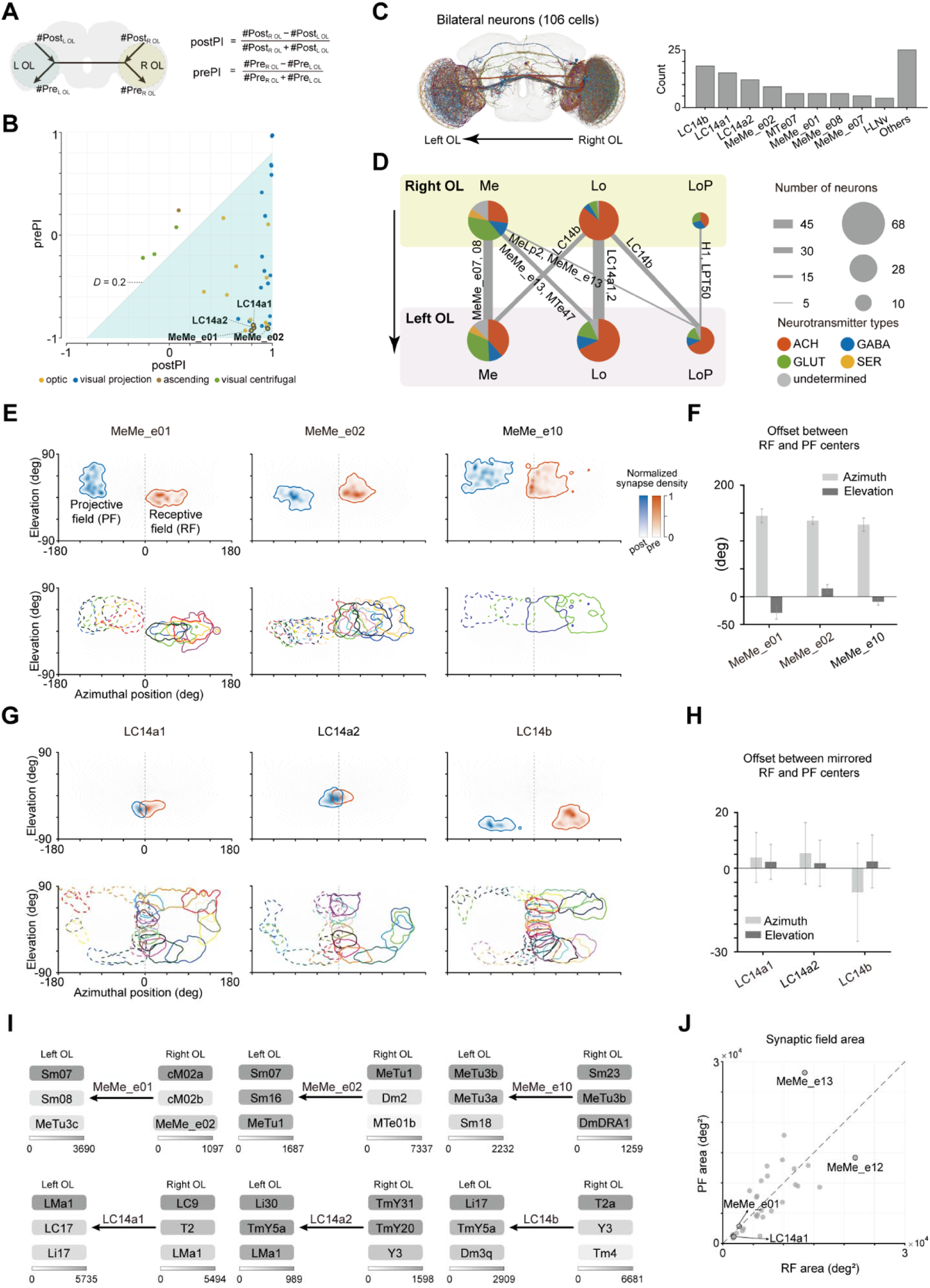
BL neurons link visual circuits in the two OLs. **(A)** Quantification of polarity indices (prePI and postPI) between the right and left OLs. **(B)** Two-dimensional scatter plot of postPI and prePI values for all cell types connecting the right OL and left OLs. Each dot represents a cell type and is color-coded by superclass. Cell types in the lower-right (cyan) transmit signals from the right OL to the left OL. **(C)** Morphologies of all BL neurons projecting from the right OL to the left OL (left), and the number of BL cells for each neuron type (right). **(D)** Schematic of neuropil-level connectivity for all BL neurons. The diameter of each circle and the width of each link are proportional to the number of cells in each group. The pie chart shows neurotransmitter identity. Cell-type names on links indicate the two neuron types with the highest synapse count for each connection. **(E)** First RF–PF pattern. Representative receptive fields (RF; red) and projective fields (PF; blue) of individual BL neurons (top), and RF (solid) and PF (dotted) contour outlines overlaid for all neurons within each BL cell type (bottom), with each neuron color-coded. **(F)** Bar plots of horizontal and vertical offsets between RF and PF centers, shown for the three BL neuron types in (E). **(G)** Second RF–PF pattern. Representative RFs (red) and PFs (blue) of individual BL neurons (top), and RF (solid) and PF (dotted) contour outlines overlaid for all neurons within each BL cell type (bottom), with each neuron color-coded. **(H)** Bar plots of horizontal and vertical offsets between mirror-transformed RF centers (reflected about the visual midline) and PF centers, shown for the three BL neuron types in (G). **(I)** Diagrams of the top three pre- and post-synaptic partners for representative BL neurons. Box color indicates the synapse count between each partner and the BL neuron (see colorbar). **(J)** Scatter plot of RF size versus PF size. A dashed line indicates the line of equality (RF size = PF size), and each dot represents a BL neuron type.

We focused our analysis on the cells with a clear directionality (*D* ≥ 0.2) for the subsequent analysis (Methods; Figure 6B). Cells with weak directionality (|D| < 0.2) were excluded as their synapses were mostly concentrated within the right OL and CB, and thus they had already been accounted for in our previous analysis. The final set of neurons connecting the two OLs is to be referred to as BL neurons hereafter (106 cells; Figure 6C, left; lower-right cyan area in Figure 6B). The most prevalent BL neuron types were LC14 (LC14b, n = 18; LC14a1, n = 15; LC14a2, n = 12), MeMe (MeMe_e02, n = 9; MeMe_e01, n = 6), and MTe07 (n = 6) (Figure 6C, right). We analyzed the input and output neuropils of the BL neurons within each OL (Figure 6D). Most innervated the same neuropils on both sides, with a few exceptions such as LC14b and MeLP2.

To assess the role of these neurons, we estimated putative receptive fields (RFs) and projective fields (PFs) from the spatial distributions of post- and presynaptic sites by mapping synapses to retinotopic columns and visual-angle space (Methods; Figures 6E-H, S5, and S6).^29,74^ This analysis revealed several distinct patterns of RF-PF combinations, two of which are described below. The first pattern comprised neurons whose RFs and PFs were separated by about 137° on average in azimuth and whose individual cells tiled the visual field with a grid of RF/PF locations that were shifted regularly along the azimuth (Figures 6E and 6F). MeMe_e01, MeMe_e02, and MeMe_e10 exemplified this organization. Notably, RFs and PFs also exhibited vertical offsets that were distinct for each cell type. The second pattern was characterized by a mirror-symmetric relationship between their RFs and PFs (Figures 6G and 6H). For example, LC14a1 and LC14a2—connectivity-defined subdivisions of LC14a^18^—formed RFs along the rim of the right visual field, whereas their PFs were positioned mirror-symmetrically across the frontal midline. LC14b, a closely related but morphologically distinct variant with an additional contralateral medulla projection,^54^ also formed a similar RF-PF relationship.

Connectivity analysis further indicated that the MeMe neurons are strongly coupled to MeTu and Sm pathways (Figure 6I, top). For example, MeMe_e10 receives prominent input from DmDRA1—a dorsal rim area (DRA) medulla interneuron implicated in skylight polarization processing^75^—and provides output to MeTu3a, MeTu3b, and Sm18 in the contralateral OL. Similarly, LC14 neurons bridge other LC neurons in both OLs by forming strong monosynaptic or disynaptic connections (Figure 6I, bottom). For example, LC14a1 integrates inputs from LC9, T2, and LMa1 and relays signals to contralateral LC neurons both directly (to LC17) and indirectly via an LMa1-to-Li17 pathway that further engages LC17, LC18, and LC12 (data not shown).

Finally, we found that the size of RF and PF varied significantly across cell types, although within a cell type, they were largely matched (Figures 6J and S6, Spearman’s ρ = 0.82, p < 0.0001). For example, the RF/PF sizes of MeMe_e12 were approximately 12 times greater than those of LC14a1 neurons. Together, our analysis highlighted the diverse patterns of interocular connections by BL projection neurons.

## Discussion

We analyzed the whole-brain *Drosophila* connectome to elucidate how visual processing emerges from a brainwide network mediated by long-range projections that link the visual system to extrinsic brain regions. By categorizing neurons according to their synapse-distribution profiles, we identified three primary functional streams: feedforward (FF), feedback (FB), and bilateral (BL). Our characterization of these pathways repositions the OL from an isolated sensory processor to an integrative hub embedded within a globally distributed computational network.

### FF bottleneck and feature reintegration

FF neurons constitute the largest class of optic-lobe-innervating projection neurons and form the primary pathway through which visual features reach the CB. Our analysis indicates that these neurons function as both a structural and functional bottleneck, transmitting visual information in a compact code shaped by extensive convergence and divergence—an architecture well suited for efficient long-range communication (Figure 2). High convergence onto FF neurons may maximize information throughput, while their distributed, overlapping outputs to postsynaptic neurons enable visual features to be reintegrated in the CB. Clustering FF neurons by their postsynaptic targets in the CB revealed functional groups that share common outputs and frequently exhibit similar visual tuning (Figure 3). This organization suggests that CB neurons typically pool matched features from distinct FF types to enhance signal robustness.^51,53^ Conversely, we also identified some instances where FF neurons with non-matching tuning converge, a configuration likely supporting the emergence of higher-order visual representations.^55–57^

### Precise recurrent loops for context-dependent visual processing

We found that the largest fraction of inputs to FB neurons originates from VPNs, and detailed connectivity analysis revealed layer-specific recurrent loops between the OL and the CB (Figure 4). Given that most FB neurons are inhibitory, these loops are well positioned to modulate the gain of specific visual features through direct inhibition or disinhibition. When recurrent circuits involve identical cell types, they may regulate feature-detection dynamics, such as adaptation via negative feedback or gain enhancement via positive feedback.^76–79,43^ In contrast, when FB neurons target visual neurons distinct from their sources of input, they may mediate mutual inhibition across parallel visual channels.^80^ Although visual input constitutes the predominant source of FB signals, a substantial fraction of FB inputs is non-visual, including signals from ascending and descending neurons (Figure 4). These inputs likely convey motor-related information previously observed across multiple OL neuropils,^11,14,81–83^ ensuring that early visual processing is continuously shaped by behavioral context. With the recent completion of whole-nervous-system connectomes,^84^ the mechanisms by which motor-related signals influence visual processing are now poised for systematic, circuit-level investigation.

### Bilateral integration and binocular vision

Most bilaterian animals possess paired visual organs, necessitating the integration of inputs from both sides to form a coherent, wide-field representation of the environment. In *Drosophila melanogaster*, the two eyes share a binocular overlap of approximately 15°, supporting the hypothesis that circuits comparing overlapping visual fields could enable depth perception via binocular disparity.^22,85^ Although LC14 neurons specifically have their receptive fields in this region, our analysis revealed that their receptive fields are not restricted to the medial rim but extend into other peripheral regions (Figure 6). Unless LC14 neurons possess specialized downstream circuits that selectively process medial rim inputs, they are unlikely to serve an exclusive role in depth perception. In contrast, MeMe neurons form a shifting azimuthal grid that likely facilitates the detection of horizon tilt (Figure 6). The strong inhibition from MeMe_e02 to MeMe_e01, combined with their reciprocal azimuthal offsets, suggests that these neurons may participate in a circuit involved in detecting horizon tilt in opposite directions. Notably, our findings on BL neurons reveal a mechanism for bilateral integration that is spatially targeted to specific visual coordinates, distinct from the global integration characteristic of wide-field optic-flow–sensitive neurons.

### From projectome to functional architecture

Vision extracts features from light impinging on its retina, which changes rapidly across space and time. To process such dynamic, high-bandwidth stimuli, the *Drosophila* brain allocates the majority of its neurons to the OL, where multiple parallel layers compute various visual features.^18,63,86^ Here, we show that visual processing is not completed within the OL but extends into the CB, where it is influenced by signals from diverse brain regions. The mammalian visual system similarly comprises an intricate network linking visual regions with the rest of the brain,^87–90^ and such projections have been thought to be directly involved in visual feature processing.^91–93^ However, prior studies were largely limited to connectivity of sparse neuron samples for their regional structures.^27,94,95^ By moving beyond these sparse, mesoscale connectivity maps, our synapse-level analysis in *Drosophila* provides a blueprint for how a compact brain achieves brainwide processing. Our findings shift the view of the *Drosophila* visual system from a FF pipeline to a distributed, recurrent engine, in which visual processing is a dynamic computation informed by visual context, bilateral input, and body state.

### Limitations of the study

Synapse-level connectomes provide unparalleled detail in wiring diagrams but lack critical physiological information—such as ion channel properties—required to fully characterize neural function. Consequently, functional inferences drawn from connectivity alone require experimental validation. Furthermore, our findings are contingent upon the FAFB-FlyWire dataset. Despite rigorous curation, these annotations may harbor residual proofreading errors or represent idiosyncratic features of a single individual rather than universal circuit motifs.^23^ Finally, as vision is inherently coupled with behavior, a holistic understanding of sensorimotor loops will require expanding these analyses to include the entire nervous system, including the ventral nerve cord.

## Supplemental Figures

**Figure S1.**
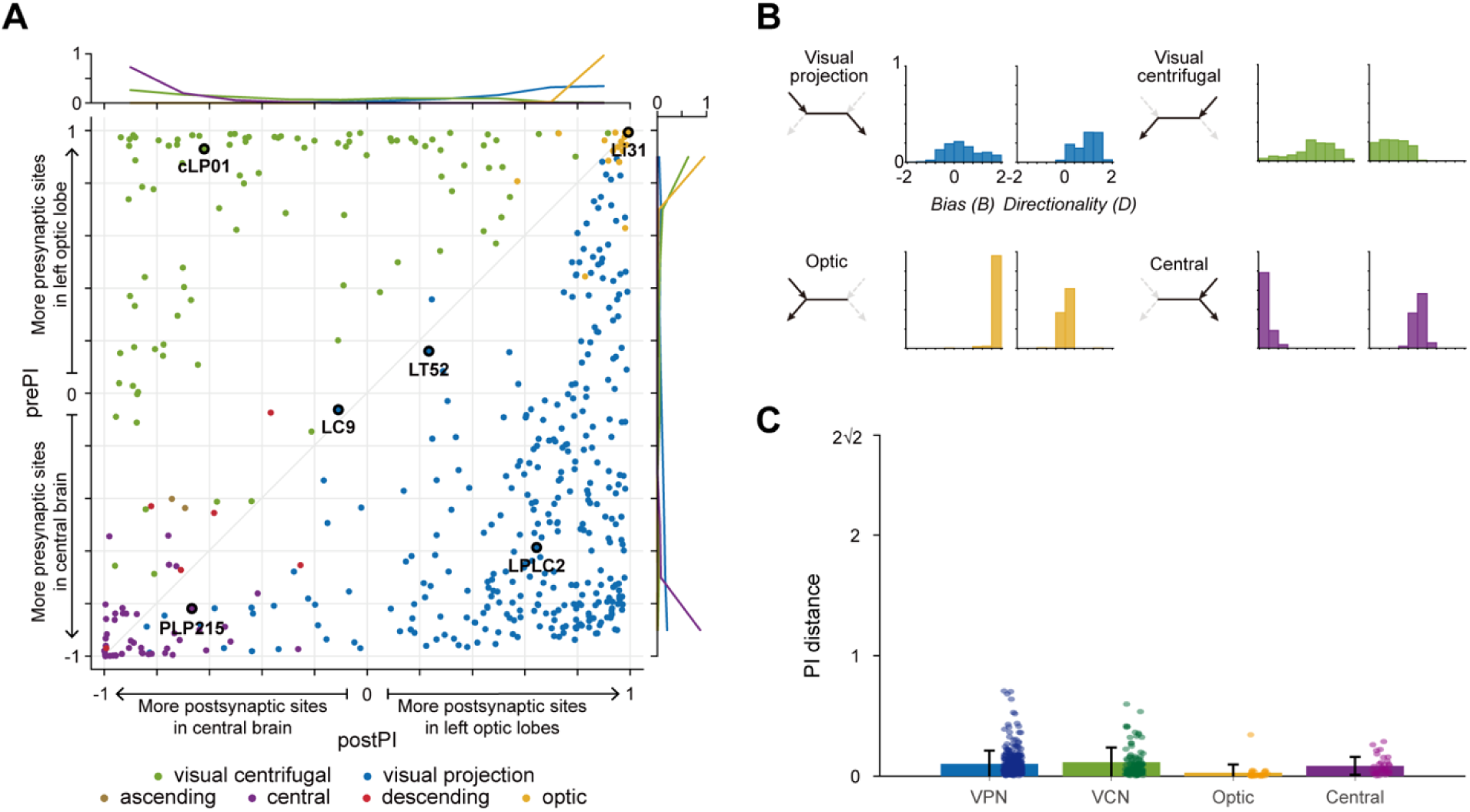
Synapse polarity-based classification of left-brain neurons. Related to Figure 1. **(A)** A two-dimensional depiction of postPI and prePI values for all cell types that connect the left OL and CB. Histograms of PI values for each cell type are shown along the top and right margins. **(B)** Histograms of spatial bias (*B*) and directionality (*D*) for left-brain neurons across four neuron classes: visual projection, visual centrifugal, optic, and central neurons. The diagram illustrating information flow is shown at the left. **(C)** Mean Euclidean distance between PI coordinates in the left and right brain for each neuron type, grouped by neuron superclass. Each dot represents the average difference in synaptic polarity across the left and right sides for a single neuron type, quantifying the bilateral consistency of our polarity estimates.

**Figure S2.**
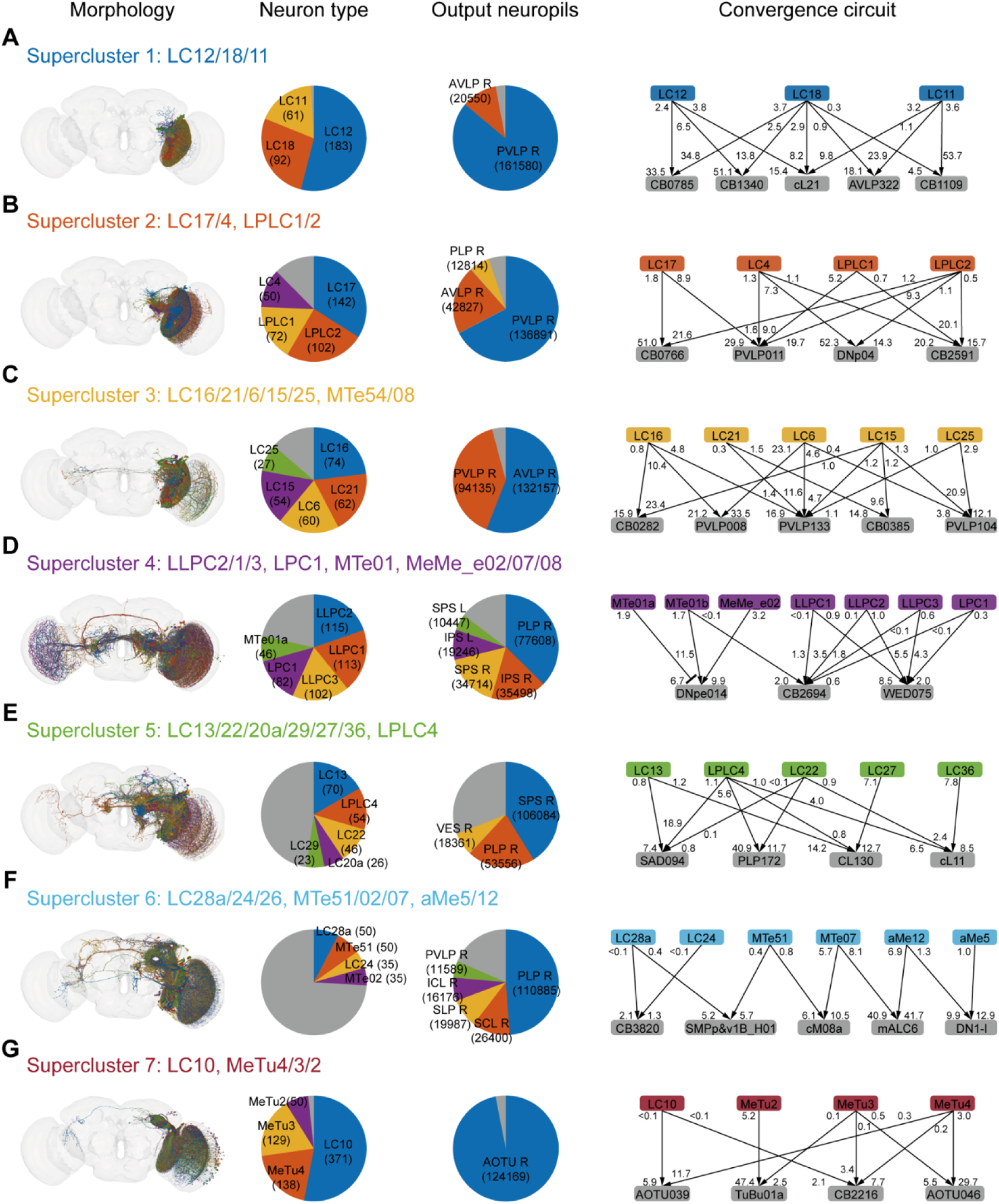
Morphologies, neuron-type composition, output neuropils, and CB convergence circuits of superclusters 1–7. Related to Figure 3. **(A–G)** Panels correspond to superclusters 1–7. Within each panel, columns (left to right) show 3D morphologies of all neurons, a pie chart of neuron-type composition (numbers indicate neuron count), a pie chart of output neuropil distribution (numbers indicate synapse count), and a schematic of representative convergence circuitry in the CB. In the convergence schematics, numbers near arrow heads indicate the fraction of the FF neuron’s total synaptic output, whereas numbers near arrow tails indicate the fraction of the CB neuron’s total synaptic input. Gray indicates categories contributing <5% (by neuron count in neuron-type pie charts; by synapse count in neuropil pie charts).

**Figure S3.**
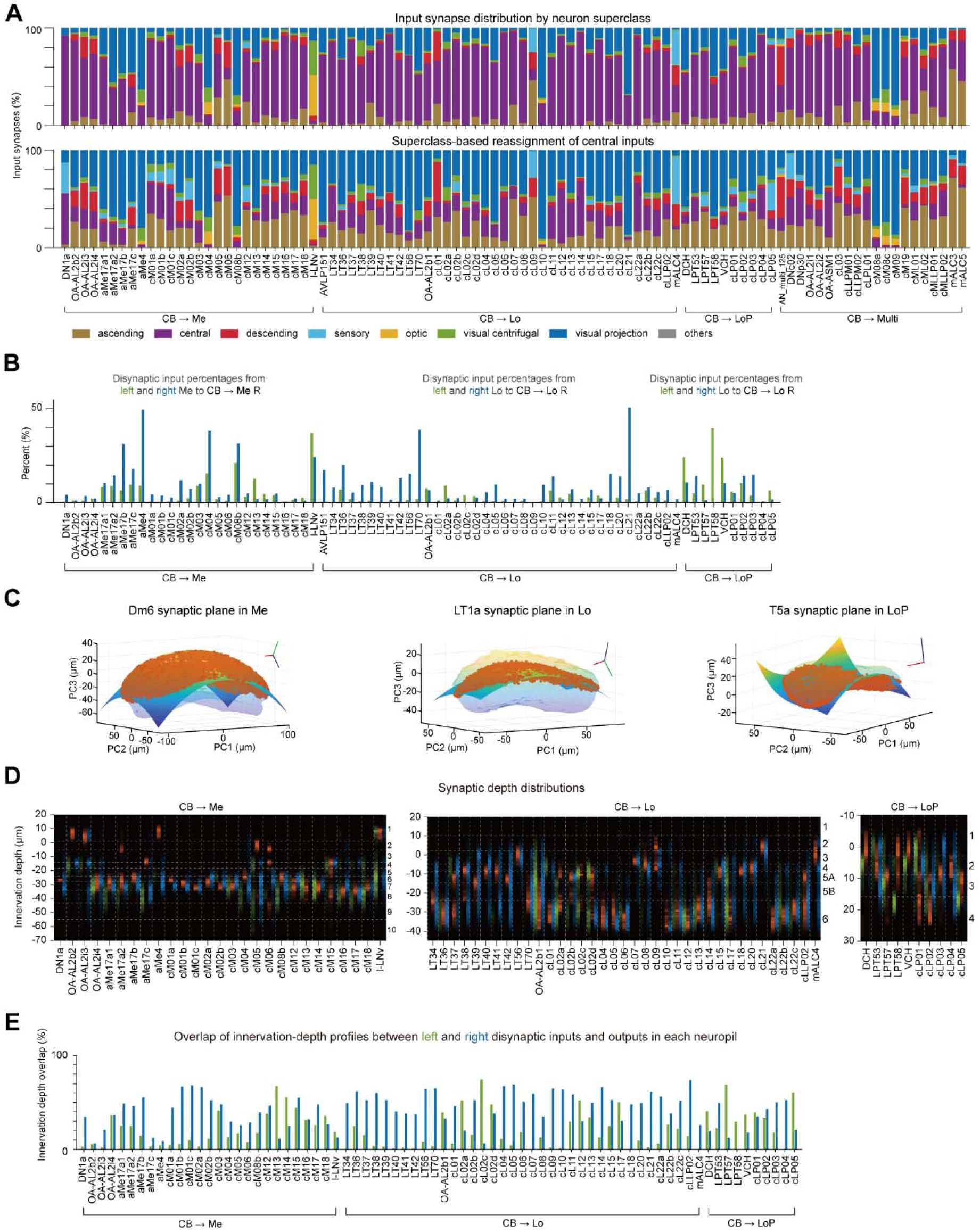
Recurrent circuits formed by FB neurons that modulate visual processing. Related to Figure 4. **(A)** Distributions of input neuron superclasses for each FB neuron group (top) and the corresponding distributions after reassigning inputs originally classified as ‘central’ to other superclasses (bottom). **(B)** Percentage of disynaptic inputs originating from the neuropil that contains each FB neuron’s presynaptic sites. **(C)** For each neuropil, synapses from a reference neuron and the layer-defining surface fitted to those synapses are shown. **(D)** Innervation-depth profiles of FB neurons for contralateral disynaptic input sites (green), presynaptic sites (red), and ipsilateral disynaptic input sites (blue). **(E)** Bar plots of overlap between innervation-depth profiles of presynaptic sites and disynaptic inputs, quantified as the summed bin-wise intersection across depth bins. Green indicates overlap with contralateral disynaptic input profiles; blue indicates overlap with ipsilateral disynaptic input profiles.

**Figure S4.**
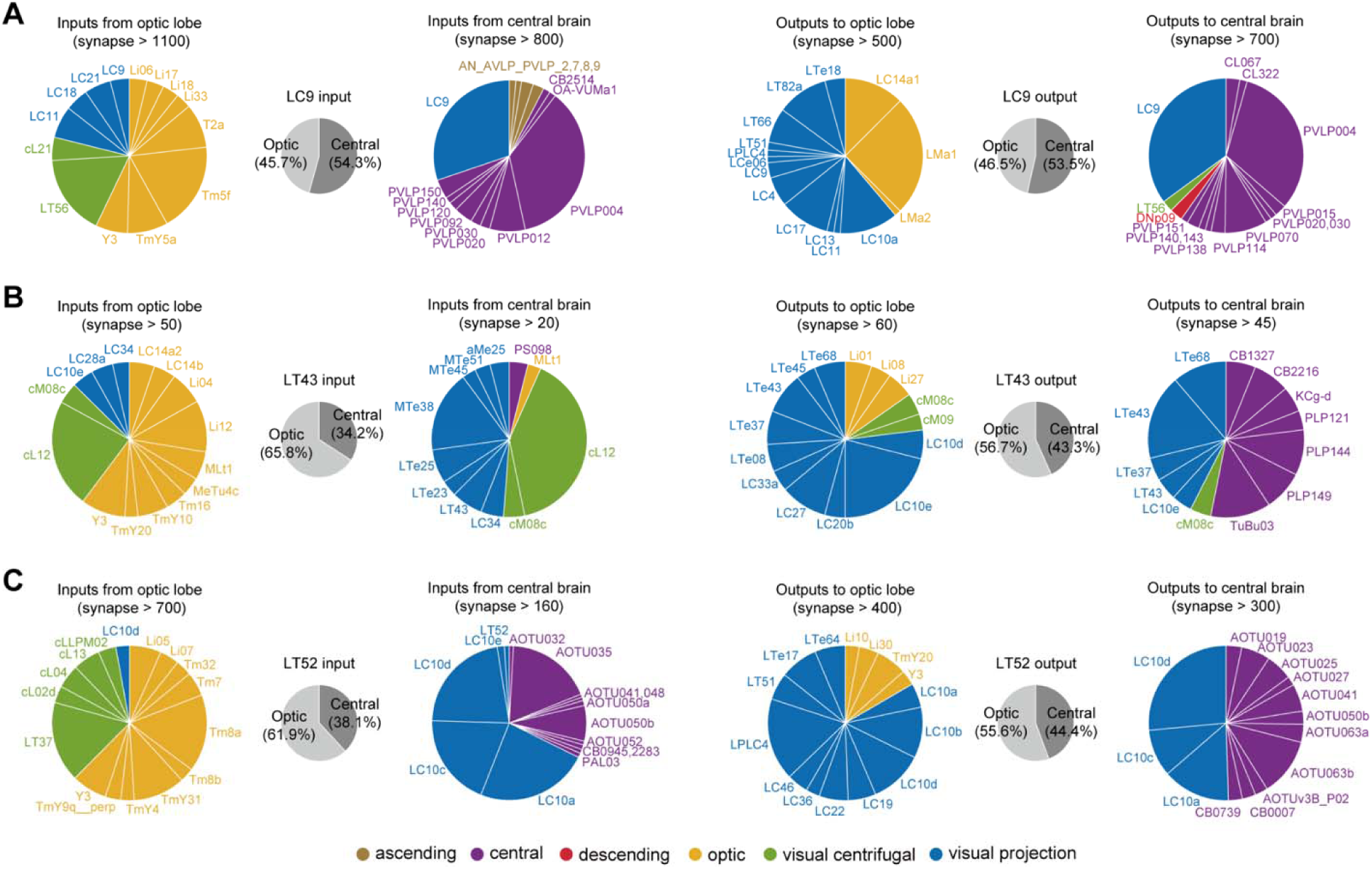
Top input/output partners of representative bidirectional neurons in the OL and CB. Related to Figure 5. **(A–C)** For LC9 (A), LT43 (B), and LT52 (C), gray pie charts indicate the proportions of total inputs and total outputs in the OL versus the CB, respectively. Partner composition is shown separately for each compartment (OL inputs, CB inputs, OL outputs, and CB outputs): cell types exceeding the synapse-count thresholds specified in each panel are displayed as pie charts, with slices normalized to the total synapses within the corresponding thresholded subset and color-coded by superclass.

**Figure S5.**
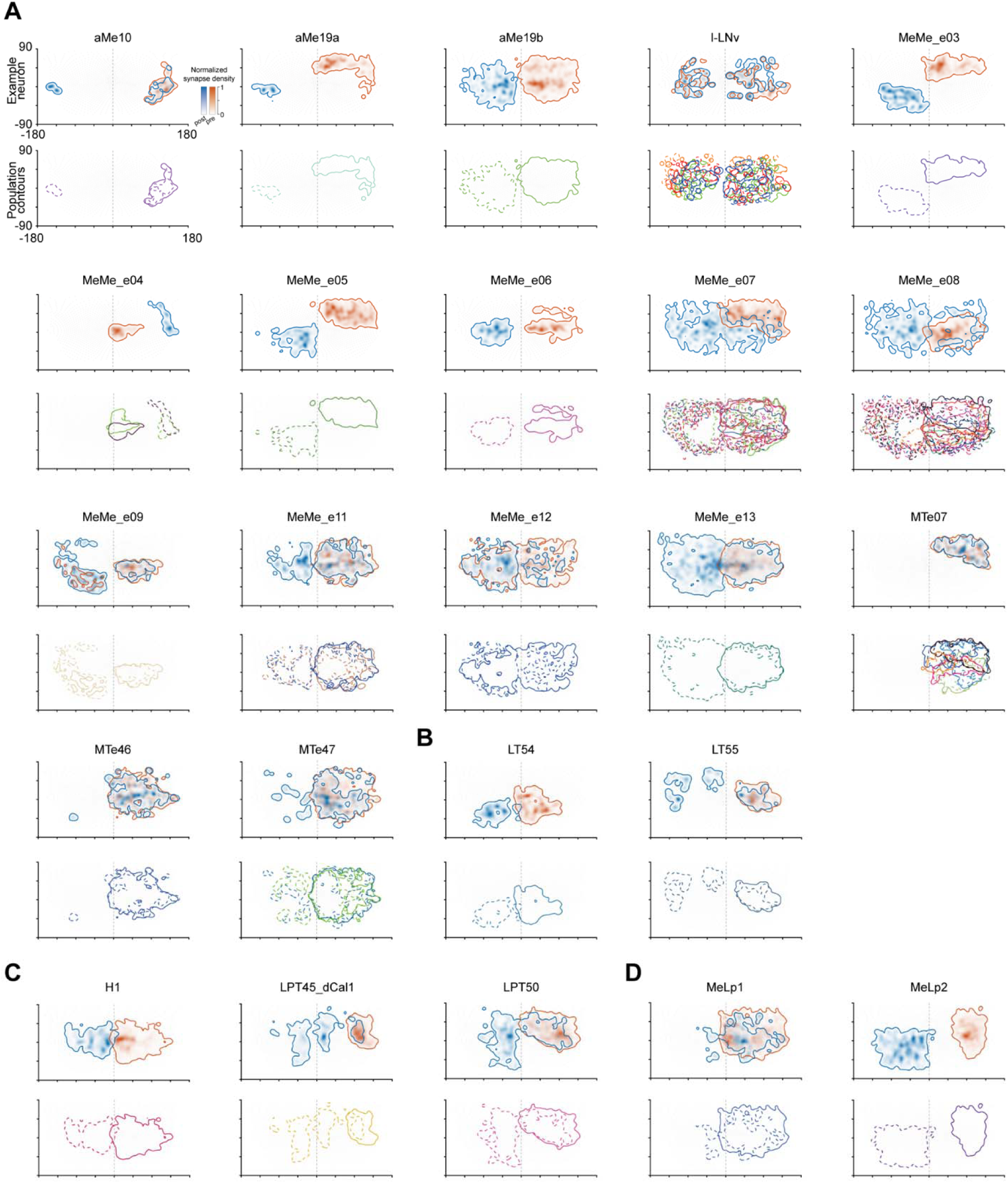
RFs and PFs of BL neurons. Related to Figure 6. **(A–D)** RFs and PFs of BL neurons, grouped by the dominant input and output neuropils: right medulla → left medulla (A), right lobula → left lobula (B), right lobula plate → left lobula plate (C), and right medulla → left lobula plate (D).

**Figure S6.**
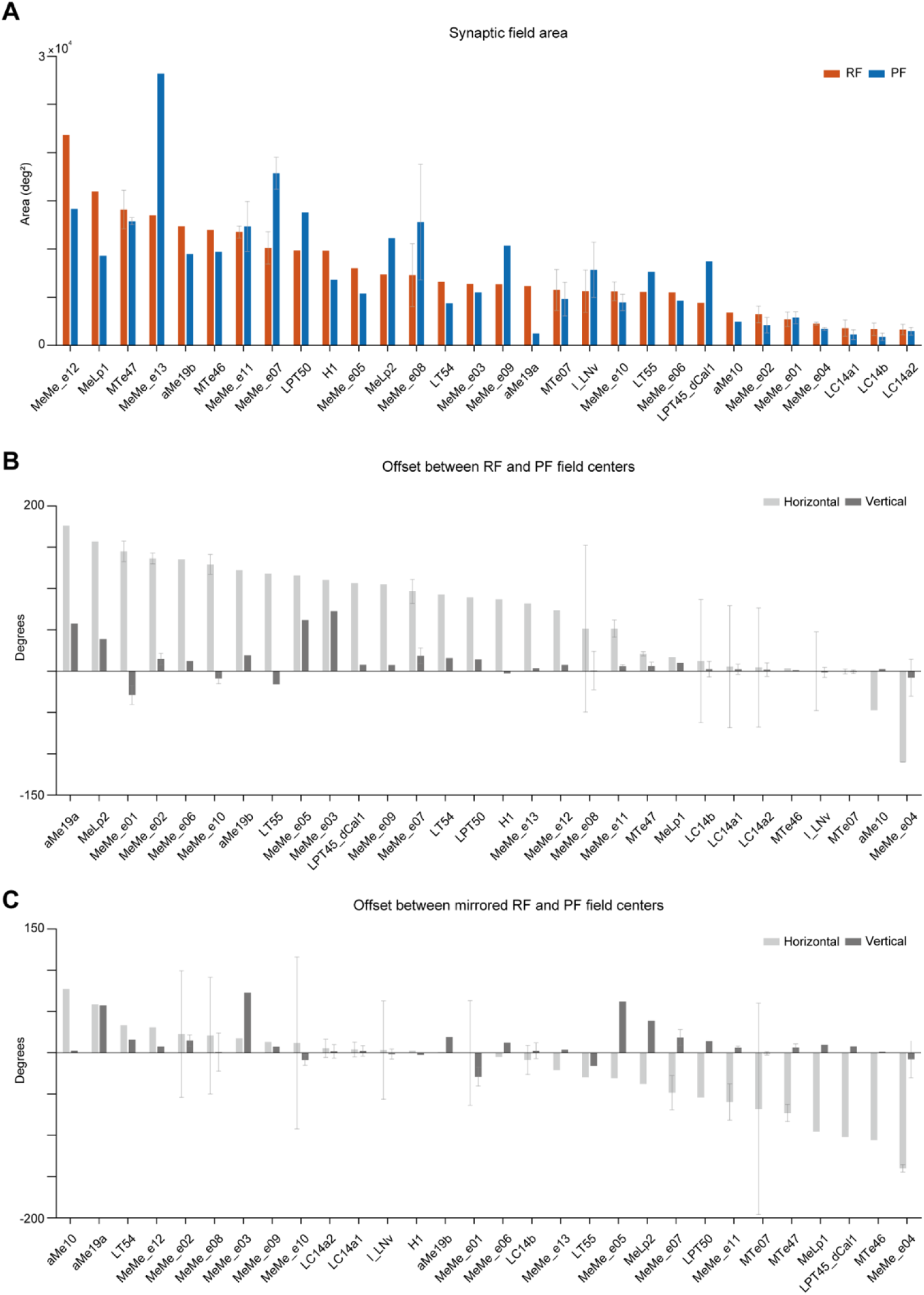
Angular size and center-offset comparisons between RFs and PFs of BL neurons. Related to Figure 6. **(A)** Bar plots of RF and PF sizes for each neuron, sorted by decreasing RF size. **(B)** Bar plots of horizontal and vertical angular offsets (degrees) between RF and PF centers, sorted by decreasing horizontal angular offset. **(C)** Same as (B), after mirroring the RF across the visual midline; angular offsets between the mirrored RF center and PF center are shown, sorted by decreasing horizontal angular offset.

## Supplemental Table

**Table S1.**
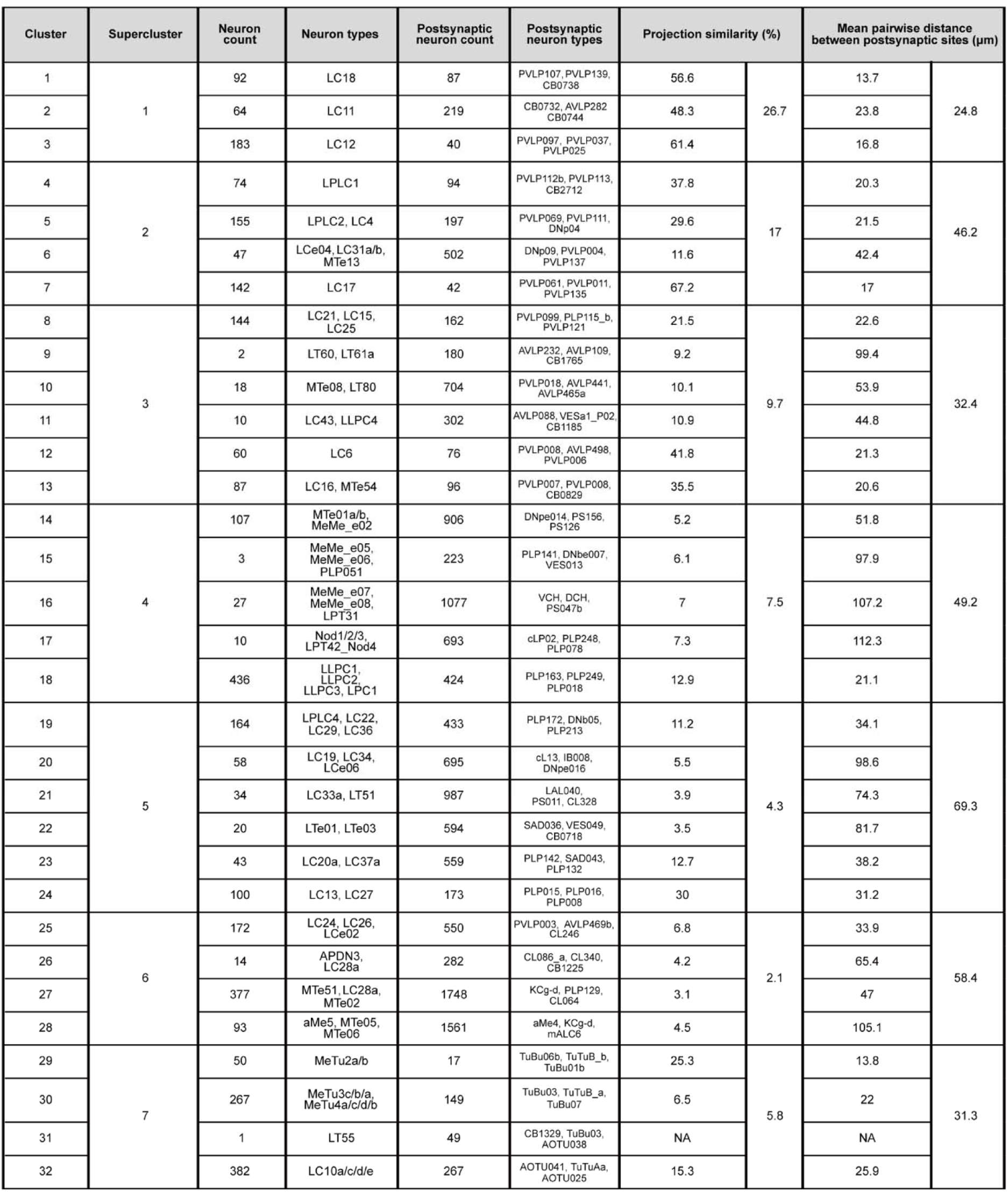
Clustering results for FF neurons based on their postsynaptic targets in the CB. Related to Figure 3. Connectivity-based clustering of FF neurons, defined by their postsynaptic target profiles in the CB, groups functionally and morphologically similar neurons into clusters and superclusters. For each FF cluster, only the three postsynaptic neuron types that receive the most synapses from that cluster are listed.

## RESOURCE AVAILABILITY

### Lead contact

Further information and requests for resources should be directed to and will be fulfilled by the Lead Contact, Anmo J Kim

### Data and code availability

Publicly available connectome data used in this study and analysis tools are listed in the key resources table. All original code to conduct these analyses will be made publicly available on GitHub upon publication. Requests for data and code prior to the publication can be directed to the authors.

## ACKNOWLEDGMENTS

The authors thank all members of the Kim laboratory for helpful discussions on the project and comments on the manuscript. We also thank Sung Soo Kim for providing resources for receptive field mapping, and the Princeton FlyWire team and consortium for the FlyWire connectome dataset and resources. This work was supported by Samsung Science and Technology Foundation (SSTF-BA2401-06) and the National Research Foundation of Korea (NRF) grant funded by the Korea government (MSIT) (RS-2022-NR070248).

## AUTHOR CONTRIBUTIONS

S.Y.K. and A.J.K. designed the study. S.Y.K. performed data analysis with inputs from A.J.K. S.Y.K. and A.J.K. wrote the manuscript.

## METHODS

### KEY RESOURCE TABLE

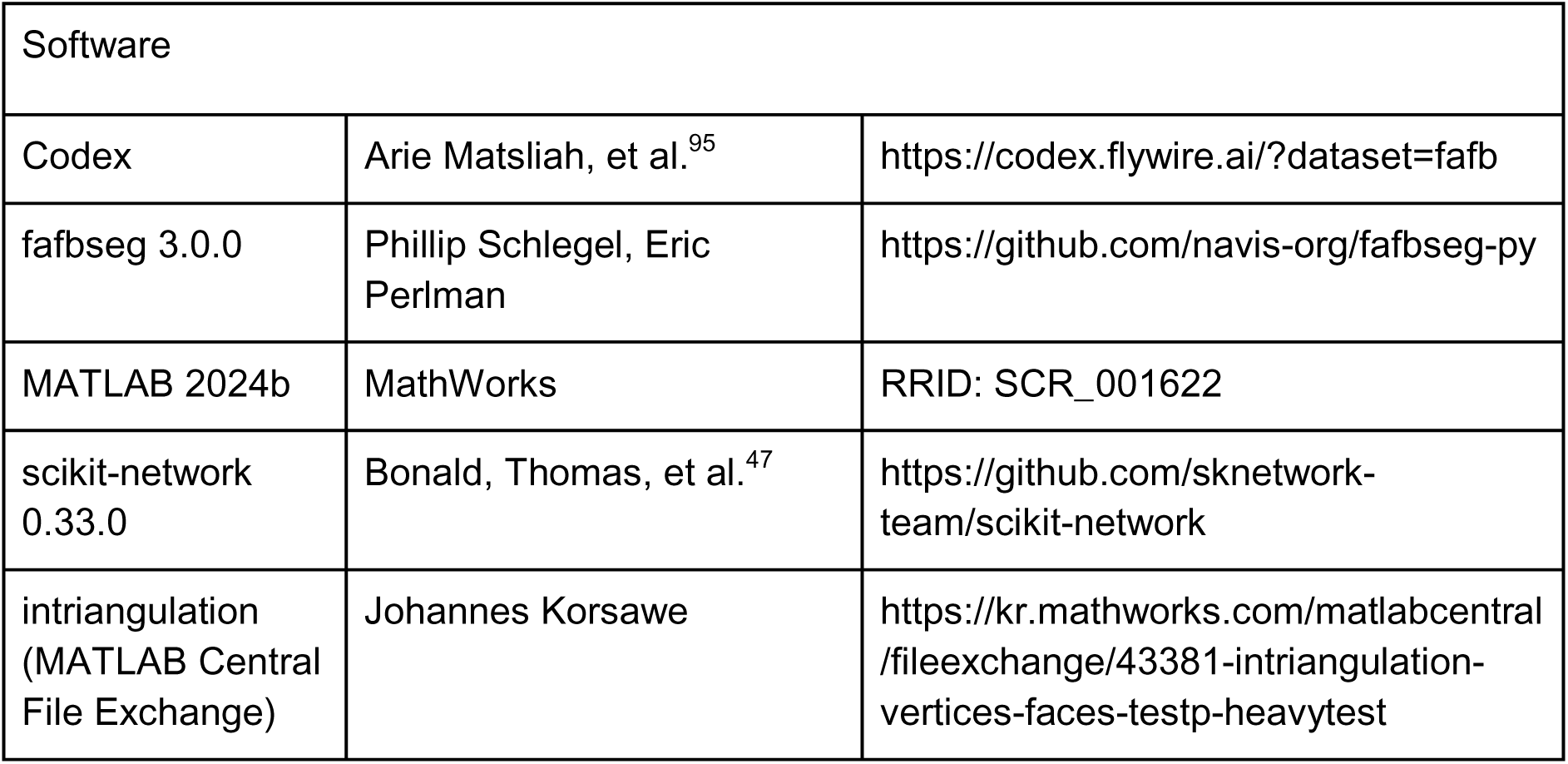

| Software |  |  |
| --- | --- | --- |
| Codex | Arie Matsliah, et al.<sup>95</sup> | https://codex.flywire.ai/?dataset=fafb |
| fafbseg 3.0.0 | Phillip Schlegel, Eric Perlman | https://github.com/navis-org/fafbseg-py |
| MATLAB 2024b | MathWorks | RRID: SCR_001622 |
| scikit-network 0.33.0 | Bonald, Thomas, et al.<sup>47</sup> | https://github.com/sknetwork-team/scikit-network |
| intriangulation (MATLAB Central File Exchange) | Johannes Korsawe | https://kr.mathworks.com/matlabcentral/fileexchange/43381-intriangulation-vertices-faces-testp-heavytest |

### EXPERIMENTAL MODEL AND STUDY PARTICIPANT DETAILS

This study did not generate new experimental animals or perform new imaging. All analyses were conducted on a previously published adult female whole-brain electron microscopy connectome of *Drosophila melanogaster,* using the FAFB-FlyWire connectome dataset (v783).^10,19,23–25^

### METHOD DETAILS

#### *Drosophila* Connectome dataset

All data analyzed in this study were derived from the *Drosophila* whole-brain connectome accessed through the FlyWire Codex browser.^95^ Neuron annotations, synapse locations, connection strengths (synapse counts), and machine-learning–predicted neurotransmitter identities were obtained from Codex and analyzed using custom scripts in MATLAB and Python. Unless otherwise stated, we included all synaptic connections and did not apply a minimum synapse-count threshold.

#### Cell filtering for the connectivity analysis (Figure 1)

We analyzed the majority of cells innervating the right OL in conjunction with the CB or left OL, excluding neurons unlikely to contribute to inter-regional visual processing based on three specific criteria. First, a neuron was included only if it possessed significant connectivity across regions, defined as having ≥ 5 postsynaptic sites in the right OL and ≥ 5 presynaptic sites in the CB, or vice versa. Second, to ensure regional specificity, neurons linking the OL and CB were excluded if more than 70% of their total input or output synapses were located outside those two regions (Figures 1–5); this served to filter out BL neurons conveying signals primarily from the left OL. Similarly, for the BL neuron analysis (Figure 6), cells were excluded if the proportion of their total synapses in the CB exceeded 70%. Finally, to maintain cell-type consistency, if more than 80% of cells within a specific FlyWire-annotated type were excluded by the aforementioned rules, the remaining cells of that type were also removed from the analysis.

#### Classification criteria for FF, FB, BD, and BL neurons (Figure 1)

To categorize neurons into functional classes, we employed specific thresholding for directionality (*D*) and bias (*B*). Neurons exhibiting clear directionality were classified as FF if *D* ≥ 0.2 and as FB if *D* ≤ -0.2. In contrast, neurons with a balanced spatial distribution and near-zero directionality (|*D|* < 0.2 and |*B|* < 1.2) were defined as BD neurons. While applying absolute thresholds for classification can be potentially misleading, these specific criteria were determined through a superclass-specific analysis of *B* and *D* indices (Figure 1E). Through this approach, we were able to effectively filter out optic and central neurons, thereby capturing an extensive population of projection neurons linking the OL and CB. The same thresholding criteria were also applied to define BL neurons.

#### Connectivity-based clustering of FF neurons (Figure 3)

To characterize how FF neurons distribute visual information within the CB, we clustered FF neurons based on their postsynaptic targeting profiles using the Leiden community-detection algorithm.^46,47^ Because this analysis focused on information transmission within the CB, we excluded (i) connections to postsynaptic neurons located in the OL and (ii) synaptic connections among FF neurons. We then constructed a weighted connectivity matrix of size 3,436 × 14,088 (rows × columns), where rows correspond to FF neurons and columns correspond to their postsynaptic partners in the CB, with entries given by synapse counts.

From this matrix, we built a bipartite graph with FF neurons as one node set and their postsynaptic target neurons as the other, with weighted edges representing synapse counts between the two sets. We then applied the Leiden algorithm using the scikit-network package.^46,47^ With the resolution parameter set to three, this procedure yielded 32 FF-neuron clusters and 32 corresponding CB-neuron clusters, where each FF cluster exhibited its strongest connectivity to its matched target-neuron partner (Figure 3C).

To determine superclusters, we quantified projection similarity between FF clusters. Specifically, we constructed a 32 x 32 matrix 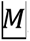, in which each element in the matrix represents the number of connections between a FF cluster and a CB cluster (Figure 3C). We then computed the FF-cluster similarity matrix as 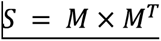, which captures similarity in postsynaptic targeting across FF clusters. The PARIS algorithm was applied to 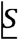 to generate a dendrogram,^47,50^ and we defined seven FF superclusters by cutting the dendrogram at a fixed threshold.

#### Output neuropil-based classification of FB neurons (Figure 4)

We classified FB neurons based on the neuropils targeted by their output synapses. For each FB neuron, we quantified the distribution of presynaptic sites across the medulla, lobula, and lobula plate; presynaptic sites in the accessory medulla were included in the medulla category. A neuron was assigned to a neuropil group if more than 10% of its total output synapses were located within that neuropil. If a neuron met this criterion for more than one neuropil, we grouped it as "Multi." Using this scheme, we identified 29 medulla-targeting FB types (CB → Me), 44 lobula-targeting FB types (CB → Lo), 10 lobula plate-targeting FB types (CB → LoP), and 18 multi-target FB types (CB → Multi) (Figure 4C and 4D). For innervation-depth calculations, we excluded multi-target FB types from the analysis.

#### Superclass-weighted reassignment of central neuron inputs to FB neurons (Figure 4C)

FB neurons receive contextual information from diverse CB populations. While some inputs originate from VPNs and ascending neurons, the majority arise from neurons categorized as central neurons (Figure 4C, top). Unlike non-central superclasses, which can be associated unambiguously with functional types (e.g., VPNs with visual signals, or ascending/descending neurons with motor signals), the roles of most central neurons are unclear. To resolve this, we reassigned the functional identity of central neurons based on their own upstream inputs. Specifically, we identified the top five cell types presynaptic to each central neuron and reassigned the synaptic count for each individual central-to-FB connection according to the proportions of those upstream types. During this calculation, we ignored any upstream central superclasses, ensuring that reassignment was performed only among functionally distinct, non-central categories (Figure 4C, bottom).

#### The proportion of disynaptic visual input to FB neurons across OL neuropils (Figure 4D)

For the four FB neuron classes, categorized by their target neuropils in the OL, we quantified the proportion of inputs originating from different OL neuropils. Specifically, we isolated presynaptic neurons that receive inputs within either OL (primarily FF neurons) and, for each, determined the proportion of total input from different OL neuropils. The synaptic weight of the connection to the FB neuron was then scaled according to these neuropil-specific proportions. For all neurons within each FB neuron type, we summed these reassigned synaptic inputs for each neuropil. Finally, we determined the proportion of the total disynaptic input originating from each OL neuropil for each FB neuron type, plotting the results as mean ± s.d. (Figure 4D). For comparison, these proportions were also calculated relative to the total synaptic input to FB neurons, including non-OL-derived signals (Figure S3B).

#### OL layer definition and innervation depth calculation (Figure 4E-G)

To measure the innervation depth of FB neurons in the OL, we defined a baseline surface for each neuropil by adapting a previously described procedure used for lobula neurons.^68^ Briefly, the workflow comprised four steps: (1) selecting layer-restricted reference neurons, (2) deriving a neuropil-aligned coordinate frame via principal component analysis (PCA), (3) constructing a baseline surface for depth calculation, and (4) computing relative synapse depth as an offset from this surface. In step 1, we chose distinct reference neurons for different neuropils: Dm6 neurons for the medulla (Layer Me1; Figure S3C, left), LT1a neurons for the lobula (Layer Lo2; Figure S3C, middle), and T5a neurons for the lobula plate (Layer LoP1; Figure S3C, right). To construct a baseline surface (step 3), we used second-order polynomials for the medulla and lobula, and third-order polynomials for the lobula plate to account for its stronger curvature. The surface fits achieved high goodness-of-fit: medulla *R*^2^ = 0.96 (right) and 0.95 (left); lobula *R*^2^ = 0.89 (both right and left); lobula plate *R*^2^ = 0.86 (right) and 0.88 (left). Finally, we defined depth ranges for each neuropil layer based on synapse-depth distributions from neuron types with established layer specificity (e.g., T4, T5, LC25, LC26, Dm6, Dm3, and LT1a), which served as empirical landmarks for defining layer boundaries (Figures 4E and S3D).

#### RF and PF estimation for BL neurons

We inferred the RFs and PFs of BL neurons from their post- and presynaptic site distributions, respectively. Because BL neurons primarily target multi-columnar interneurons, traditional connectivity-based mapping^29,96^—which relies on single-column partners like Mi1—is insufficient. We therefore developed a synapse-based mapping approach in three steps.

First, for the medulla, we utilized a published microCT-based eye map that assigns each Mi1 column to specific azimuthal and elevational coordinates in the visual field.^29,74^ For the lobula and lobula plate, we mapped columns to a visual area based on their connectivity to the medulla. Specifically, for each column-defining cell, such as Tm3 for the lobula or T4a for the lobula plate, we identified upstream Mi1 neurons and calculated a weighted visual field using normalized Mi1→Tm3/T4a synapse counts.

Second, we projected each BL synapse onto a neuropil-specific baseline surface derived from layer-restricted landmark neurons. Column centers were established by projecting and averaging the presynaptic sites of these landmark neurons onto the same surface. Projections were orthogonal for the medulla and lobula, whereas the PC3 axis was used for the lobula plate to account for its complex curvature. Each BL synapse was then assigned to the nearest projected column center.

Finally, RF and PF maps were generated by aggregating synapses into 2D heatmaps with 2° bins based on their assigned visual coordinates. To minimize sampling noise and approximate ommatidial optical blur, maps were smoothed with a Gaussian filter (FWHM=9.5)^97^ using circular wrap-around in azimuth and boundary padding in elevation. Finally, maps were peak-normalized, and the 10% isodensity contour was extracted for comparison across neurons.

### QUANTIFICATION AND STATISTICAL ANALYSIS

#### Data processing and reporting conventions

All analyses were performed in MATLAB and Python. Metrics were initially computed for individual neurons and subsequently averaged for visualization at the cell-type level, with the following exceptions: the marginal histograms in Figures 1D and S1A, the histograms in Figures 1E and S1B, the FF neuron clustering analyses in Figure 3, and the representative BD neuron analyses in Figure 5. For box-and-whisker plots, the center line indicates the median, the box spans the interquartile range (IQR; Q1–Q3), and whiskers extend to 1.5 x interquartile range. Unless otherwise stated, data are reported as mean ± s.d.

#### Histograms of synaptic coordinate distributions (Figures 1B and 1H)

To compare the spatial distributions of synaptic inputs and outputs, we constructed overlapping histograms along the axis of interest. For each neuron, we first identified the minimum and maximum synapse coordinates along that axis and used their difference to define the axis range. We then divided this range into 100 equally spaced bins, and assigned each synapse to a bin based on its coordinate.

#### Statistical analysis and significance criteria

Statistical significance was assessed using Mann–Whitney U tests in Figures 2C–E, 3F–G, and 5C–E and 5G, as indicated in the figure legends. For Figures 2C–E and 5C–E and 5G, results were considered significant at a Bonferroni-corrected threshold of *p* < 0.0167. For Figures 3F–G, *p* values are reported without Bonferroni correction.

#### Intra-type clustering consistency (Figure 3D)

To quantify the consistency with which individual neurons of the same type were assigned to the same cluster, we computed the fraction of all neuron pairs within that type that were co-assigned to the same cluster. We termed this metric “intra-type clustering consistency.”

#### Synapse-weighted Jaccard index for projection similarity and reciprocity

Projection similarity in Figure 3E and the proportion of reciprocal connections in Figures 5F and 5G were quantified using the synapse-weighted Jaccard index. For two connectomic profiles, *A* and *B,* defined over a neuron index i––with weights a_i_ and b_i_ representing synapse counts––the index was computed as: *J_W_(A, B)* = ∑_i_ min(*a_i_, b*_i_ ) / ∑_i_ max (*a_i_, b_i_*). *J_W_* ranges from 0 (no overlap) to 1 (identical weighted distributions). In Figure 3E, we used this metric to compute the similarity between the CB target profiles of two neurons. To quantify reciprocity in Figures 5F and 5G, we calculated the Jaccard index between an individual neuron’s upstream and downstream partner profiles. In all cases, weights were defined by cell-to-cell synapse counts.

#### Pairwise LC9 distance within the OL (Figure 5)

Pairwise LC9 distance within the OL (Figures 5I and 5J) was computed as the mean Euclidean distance between the synapse-location distributions of two LC9 neurons. This synapse-based distance metric served as a proxy for the spatial separation between their neuritic arbors in the OL.

#### Area and centroid quantification of RF and PF

We quantified the area and centroid of both RFs and PFs from normalized heatmaps using a single contour level (0.1). We defined the active region as the set of bins with normalized intensity ≥ 0.1, and computed its area by counting these bins and multiplying by the bin area (2° × 2° = 4 deg²). The center was computed as an intensity-weighted center within the active region rather than as a geometric centroid. For latitude (θ), we calculated the weighted arithmetic mean of latitude values using heatmap intensity as the weight. For longitude (φ), which is circular (−180° and 180° are equivalent), we computed a circular weighted mean by converting each longitude to a unit-vector representation, summing these vectors weighted by heatmap intensity, and converting the resulting vector back to an angle via the four-quadrant inverse tangent function.

## Declaration of generative AI and AI-assisted technologies in the manuscript preparation process

During the preparation of this work, the authors used ChatGPT and Google Gemini to edit and improve the draft’s clarity, and to assist with writing analysis and visualization code. After using these tools, the authors reviewed and edited the content and take full responsibility for the content of the published article.

